# Complementary ecomorphological patterns in brain shape and size evolution across the avian radiation

**DOI:** 10.1101/2025.10.14.682392

**Authors:** Talia Lowi-Merri, Akinobu Watanabe

## Abstract

Like mammals, birds have exceptionally large brains for their body sizes, which is thought to have facilitated their diversification through environmental change, and has been associated with many ecological adaptations. Although size metrics are informative and well-studied, they are limited in their ability to characterize localized neuroanatomical variation. Size and shape together can holistically capture the diversification of avian brain form, and how these changes relate to function and ecology. We employ high-density geometric morphometric (GM) shape data across 300 avian species to capture shape variation in whole brain endocasts, and compare how evolutionary patterns differ between shape and size. While ecomorphological signal is mixed across brain regions, we find that the majority of shape variation falls along an ‘elongated- globular’ spectrum, where terrestrial and aquatic birds generally have anteroposteriorly elongated brains, while aerial bird brains are typically more globular. The supermajority (>90%) of endocranial shape variation is independent of evolutionary allometry, demonstrating that shape characterizes key aspects of avian neuroanatomy that are missing from volumetric data. Our results suggest that neuroanatomical diversity in birds is not driven by the dominance of any particular factors, but rather through clade-specific divergence in localized brain structures from the avian-wide allometric and ecomorphological patterns.

## INTRODUCTION

The evolutionary transition from non-avian dinosaurs to crown birds (Aves; hereby ‘birds’) marks a major transformation in vertebrate evolution, giving rise to one of the most diverse groups of terrestrial vertebrates following a drastic change in body plan ^1–3^. One such change was the evolution of a highly encephalized brain, which may have contributed to the survivorship of crown bird lineages across the end-Cretaceous mass extinction event, and the ability to diversify and adapt to different environments^4–7^. These exceptionally large brains, especially in crows and parrots ^8^, are often accompanied by unique cognitive abilities such as innovativeness and tool- use ^9–13^, behavioural flexibility to environmental variation ^4,14–16^, greater ‘social intelligence’ characterized by strong social bonds ^17–19^, and song-learning shaped by neurons that resemble those of language-learning in humans ^20,21^. As such, the avian brain provides a compelling comparative system to mammals for the evolution of encephalized brains capable of higher cognitive functions.

Previous macroevolutionary studies have largely relied on volumetric data to investigate evolutionary trends and ecological correlates^4,5,16,22^. For example, initial pulses of encephalization occurred prior to and within crown birds. These trends were due, in part, as a result of body size reduction occurring more rapidly than brain size reduction along the dinosaur- bird lineage, which presumably relates to reducing body mass for flight or gliding ^5^. Additionally, ecological traits that reflect sensory abilities, such as vision and spatial orientation, are strongly correlated with both global and regional brain size metrics ^23–26^. For testing functional and ecological correlates, these studies on volumetric data often follow the ‘principle of proper mass’^27^, which in an evolutionary perspective, implies that relatively larger brain structures reflect greater demand and selection for the functions associated with those regions. This holds true in several aspects of the avian brain: an enlarged Wulst is correlated with binocular depth perception ^28^, and an enlarged medial spiriform nucleus connecting the associative pallium with the cerebellum is correlated with complex foraging behaviours ^29^. While size measurements are informative and have led to important insights on brain evolution, they are limited in their ability to characterize brain morphology. For instance, brains with equivalent relative and absolute sizes could still diverge in other morphological aspects, such as interregional connectivity and configuration, especially in macroevolutionary studies that encompass large interspecific variation^30^. Further, functional ability is often associated with structures across multiple regions, or manifests as subtle, localized differences within a region (e.g., visual Wulst on the dorsal aspect of the cerebrum). One aspect of neuroanatomical variation that is not captured by size is the spectrum from elongated to rounded, or ‘globular’, brain shapes^31–35^. Globularization is characterized by a restructuring of brain tissue configuration in physical space, such as the centralization of the thalamus^34,35^. This anatomical rearrangement is sometimes explained through the ‘spatial packing’ hypothesis^36^, in which large brains, and greater quantities of brain tissue, are accommodated within the cranial cavity by becoming more rounded in shape and through flexion of the cranial base^37^. Spatial packing is generally associated with large brains, however the degree to which globularization is correlated with brain or body size, and whether additional ecological or evolutionary variables are associated with this axis of variation, is unclear^33,34^. Examining shape variation allows us to cohesively capture 1) the proportional size differences of brain regions relative to one another, 2) subtle, localized differences within major regions that may be functionally relevant (e.g., the visual Wulst), and 3) relative positions (i.e. configuration) of brain regions. While extensive 3D characterization and ecomorphological analyses have proceeded in avian skeletal systems^38–40^, investigations into brain and endocranial shape variation have been limited to either relatively simple shape characterization with few landmarks ^38,41,42^ or have been applied to a sparse taxonomic sampling given the speciose avian diversity ^26,43–45^. This leaves a major gap in our understanding of the ecological, evolutionary, and anatomical correlates of avian brain shape.

Applying a series of phylogenetically-informed methods on 300 crown bird species and six non-avian dinosaurs, we employ a high-density geometric morphometric (GM) approach to examine the macroevolutionary and ecomorphological patterns in avian brain shape, using digital endocasts (internal mold of brain cavity). While internal brain anatomy is difficult to study with a broad phylogenetic scope and in fossil specimens, we provide a comprehensive analysis of the surface anatomy known to accurately represent external brain shape in birds and paravian dinosaurs^43,44,46,47^ and also correlate with some of the functionally relevant internal brain regions^46^. Whole endocast and regional shapes are compared with a wide breadth of ecological variables, including diet, foraging behaviors, locomotor lifestyle, and developmental mode (altriciality and precociality), to explore how ecological factors are associated with brain shape. Further, quantifying rates of neuroanatomical shape evolution can clarify if evolutionary shifts correspond with major transitions in ecological niche. To investigate how both brain size and shape provide complementary insights, we conducted analyses on both modalities of neuroanatomical characterization. With these data, we examine 1) major axes of brain shape variation in extant birds using principal component analysis (PCA); 2) rates of brain shape evolution across the avian phylogeny; 3) link between brain shape and size evolution through combined analysis of geometric morphometric and size data; 4) ecomorphological signal in brain morphology, including associations with diet, developmental mode, and locomotion.

### Institutional abbreviations

**FMNH,** Field Museum of Natural History, Chicago, IL, USA; **IGM,** Institute of Geology, Mongolian Academy of Sciences, Ulaanbaatar, Mongolia; **IVPP,** Institute of Vertebrate Paleontology and Paleoanthropology, Beijing, China; **NHMUK PV OR**, Natural History Museum, Palaeontology Vertebrates, Ornithology, London, UK.

## RESULTS

### Endocranial shape variation

To visualize major aspects of variation in endocranial shape, we produced a phylomorphospace through PCA on GM data (Fig. 1). In the dataset comprising extant birds and non-avian coelurosaurian dinosaurs, the first two principal components (PCs) accounted for 55.9% of the cumulative shape variance. Subsequent axes each accounted for < 10% of the shape variance and thus will not be discussed, but are included in comparative shape analyses. PC1 (30.9% of variation with fossils, Fig. 1; 30.1% without fossils, Fig. S1, S2) clearly separates the non-avian dinosaurs from crown birds, as shown in previous studies ^43,45,48^, accounting for the relative size and anterodorsal position of the cerebrum to the other brain regions. Positive PC1 values are associated with a proportionately larger, dorsally positioned cerebrum and an anteroposteriorly short cerebellum. Taxa with the most extreme positive PC1 values include passerine birds such as corvids, furnariids *Scytalopus* and *Campyloramphus*, as well as the soaring parrot *Nestor* and the kiwi *Apteryx*. Negative PC1 values are associated with a relatively smaller, rounder, and anteriorly positioned cerebrum, an anteroposteriorly longer cerebellum and a dorsoventrally longer optic lobe. Extreme negative PC1 values are occupied by the fossil taxa, aquatic pursuit hunters like the Sulidae (anhingas and cormorants), Galliformes (landfowl), and Caprimulgiformes (nightjars). PC2 (25.04% of variation with fossils; 25.3% without fossils) is characterized by degree of dorsoventral flexion of the brain, especially the cephalic flexion between the cerebrum and midbrain. Positive PC2 values correspond with a dorsoventrally curved, anteroposteriorly compressed cerebrum, a ventrally angled cerebellum and brainstem, and a dorsoventrally longer optic lobe. The most extreme positive PC2 values are occupied by both apodiform groups, the hummingbird *Archilochus* (nectarivore) and the swift *Hemiprocne* (aerial insectivore), which are also clustered by diet. Negative PC2 values are associated with more elongated brains, with a dorsoventrally flatter cerebrum, cerebellum, and brainstem, with extreme values being occupied by *Tyrannosaurus rex*, the most basally divergent fossil in the dataset, and the aquatic predators *Anhinga* and *Phalacrocorax*.The phylomorphospace shows some phylogenetic structure in morphological variation (K_mult_ = 0.62, *p* = 0.001), most clearly with Apodiformes. Both the whole endocast and regional datasets showed statistically significant levels of phylogenetic signal (Table S1). These analyses demonstrate that the major axis of endocranial shape variation is along the globular-elongated spectrum, an aspect of morphology that cannot be captured by volumetric data alone.

**Fig. 1:**
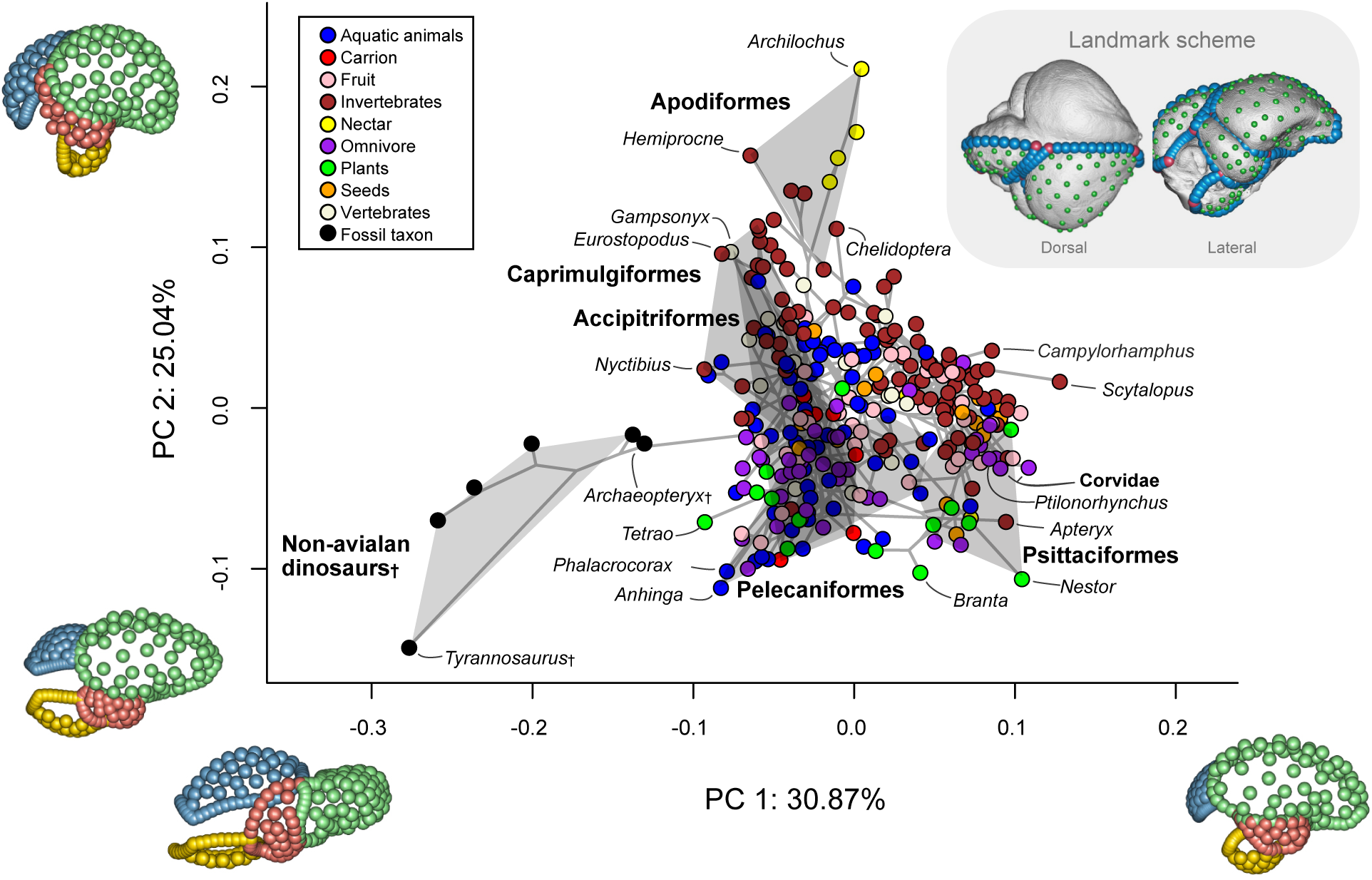
Phylomorphospace of the first two principal component (PC) axes highlighting the major aspects of endocranial shape variation in the extant and fossil dataset. Points are colored by discrete dietary category. Inset images within the plot illustrates the landmark scheme used in this study on the endocast of *Picoides pubescens* (FMNH 290218). Inset images along the axes depict shape changes associated with PC1 and PC2 axes, colored by region (green; cerebrum; red: optic lobe; blue: cerebellum; yellow: brainstem)

### Evolutionary rates of endocranial shape

Examining rates of neuroanatomical evolution in the avian brain can allow us to identify whether rates change in concert with major ecological transitions in birds. To assess the evolutionary tempo and mode of brain shape, we fit multiple variable-rates models of evolution to the extant + fossil shape dataset. These tests used PC axes accounting for 99% of shape variance to analyze rates of shape change per million years, and endocast shape was analyzed both globally and regionally. A Brownian motion model of trait evolution with a λ transformation, in which λ = 0 indicates that there is no phylogenetic signal in the trait and λ = 1 indicates the trait evolving as expected under Brownian motion^49^ was the best-supported evolutionary model for each brain partition (total endocast: λ = 0.6; cerebrum: λ = 0.61; optic lobe: λ = 0.69; cerebellum: λ = 0.47; brainstem: λ = 0.58). We then mapped these rates of morphological change onto time-calibrated phylogenies to visualize how evolutionary rates differ across taxonomic groups (Fig. 3; Fig. S3). Elevated rates are concentrated at the base of the tree among non-avian dinosaurs for all brain partitions. For the whole endocast (76 PC axes = 99% of variation) and within crown birds (Fig. S3), high rates are observed in the branches leading to *Apteryx* (kiwi), Anserinae (swans, geese), Apodiformes (nectarivorous hummingbirds and aerial insectivorous swifts), Rallidae (rails), independently across several diving birds, Aegypiinae (Old World vultures), Psittaciformes (parrots), and corvids (crows, ravens). Evolution of the cerebrum (65 PC axes) also shows elevated rates within branches of Psittaciformes and within Sulidae and its phylogenetically and ecologically close group Threskiornithidae (ibises, spoonbills), as well as in the branch leading to *Apteryx* (Fig. 3A). Evolution of the optic lobe (37 PC axes) was more variable throughout the tree, with high rates seen at the base of Apodiformes, the charadriiform clade containing Stercorariidae + Alcidae (auks and jaegers), within Sphenisciformes (penguins), and within Accipitrimorphae (aerial vertebrate predatory hawks, eagles, and vultures), and the branch leading to Passeriformes, the hyperdiverse clade of perching birds (Fig. 3B). Evolutionary rates remained generally low among crown birds for the cerebellum (46 PC axes) relative to lineage leading to non-avian dinosaurs, with higher rates observed at the base of Anatidae (ducks, geese and swans) and at the base of the passerine family Ptilonorhynchidae (bowerbirds) (Fig. 3C). The brainstem (40 PC axes) showed elevated rates at the base of Caprimulgiformes (nightjars) as well as throughout Apodiformes, the Piciformes suborder Galbuli (jacamars and puffbirds), and especially accelerated rates at the base of Corvidae (Fig. 3D). Generally, substantial rate changes occur during shifts into new ecological niches, especially during the transition to hyperaeriality in Apodiformes across all brain regions.

### Endocranial size and shape allometry

To test for the relationship between brain and body size, we regressed log-transformed endocast centroid size, a proxy for brain size, against log-transformed cube-rooted body mass. As expected, endocast size was strongly correlated with body mass (R^2^ = 0.89, slope = 0.62, p < 0.005) (Fig. 2), with Palaeognathae (e.g., *Eudromia, Struthio, Rhea*) and Galliformes (e.g., *Tetrao, Gallus, Bonasa)* exhibiting smaller endocast sizes relative to body than expected, while Psittaciformes (e.g., *Nestor, Psittacus, Deroptyus*) exhibited larger endocast sizes than the avian- wide trend. Allometric scaling of shape was visualized by plotting the regression score representing overall shape against log-transformed centroid size and body size (Fig. 2B–D). Endocast shape showed statistically significant correlation with both size metrics, with similar coefficients of variation with brain centroid size (R^2^ = 0.064, p = 0.001) and body size (R^2^ = 0.063, p = 0.001). This result indicates that size accounts for proportionately small amount of total shape variation, supported visually by the substantial scatter surrounding the main trend and the principal shape variation associated with globular-elongated morphologies (Fig. 2).

**Fig. 2:**
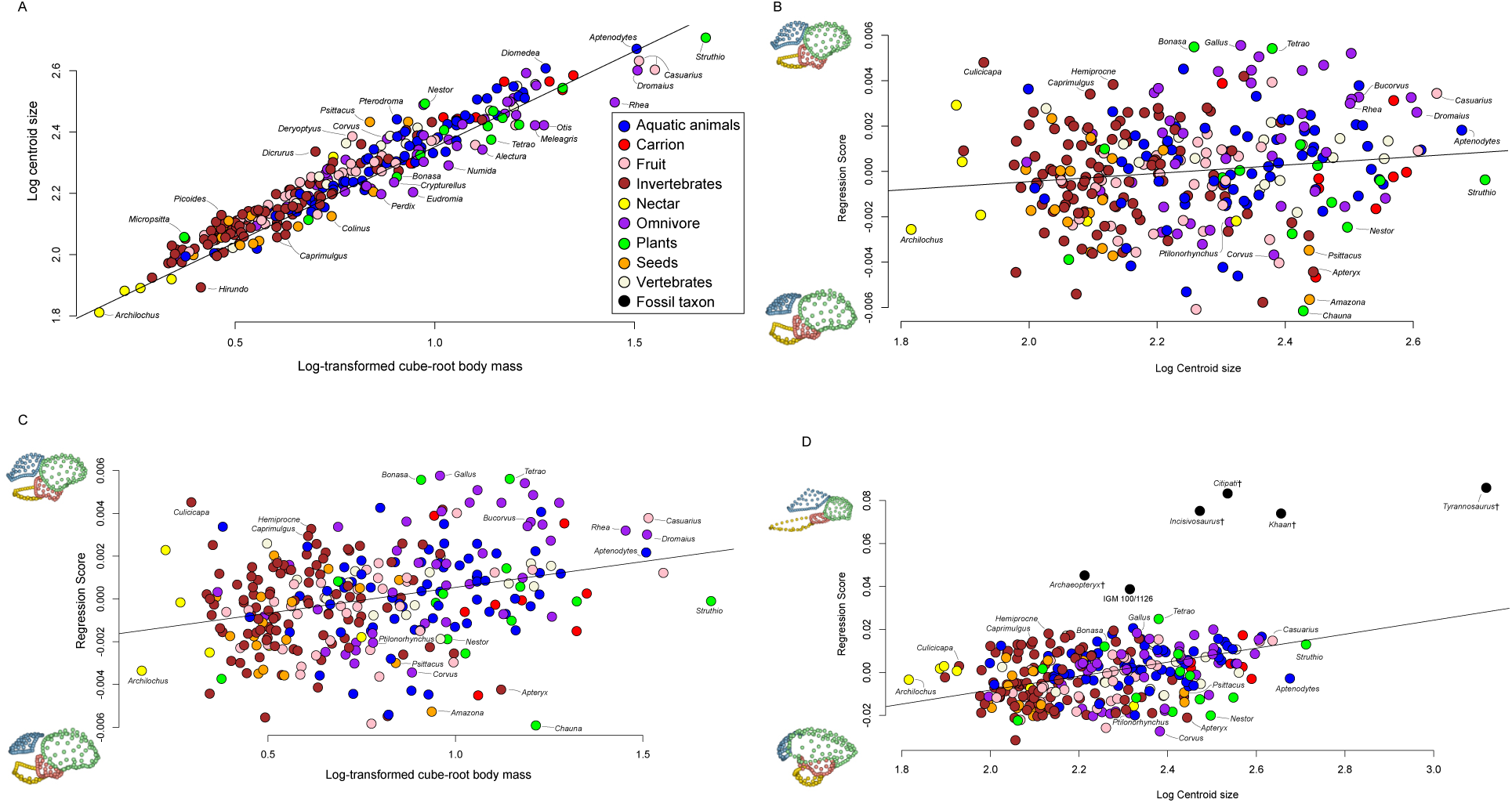
Plots of shape variable against size metrics colored by discrete dietary categories. A. Log-transformed cube-rooted body mass plotted against log-transformed centroid size. B. Log centroid size against regression score, a metric representing overall shape captured by a PGLS regression, of the extant-only dataset. C. Log- transformed cube-rooted body mass against regression score. D. Log centroid size against regression score of the dataset including both extant and fossil taxa.

**Fig. 3:**
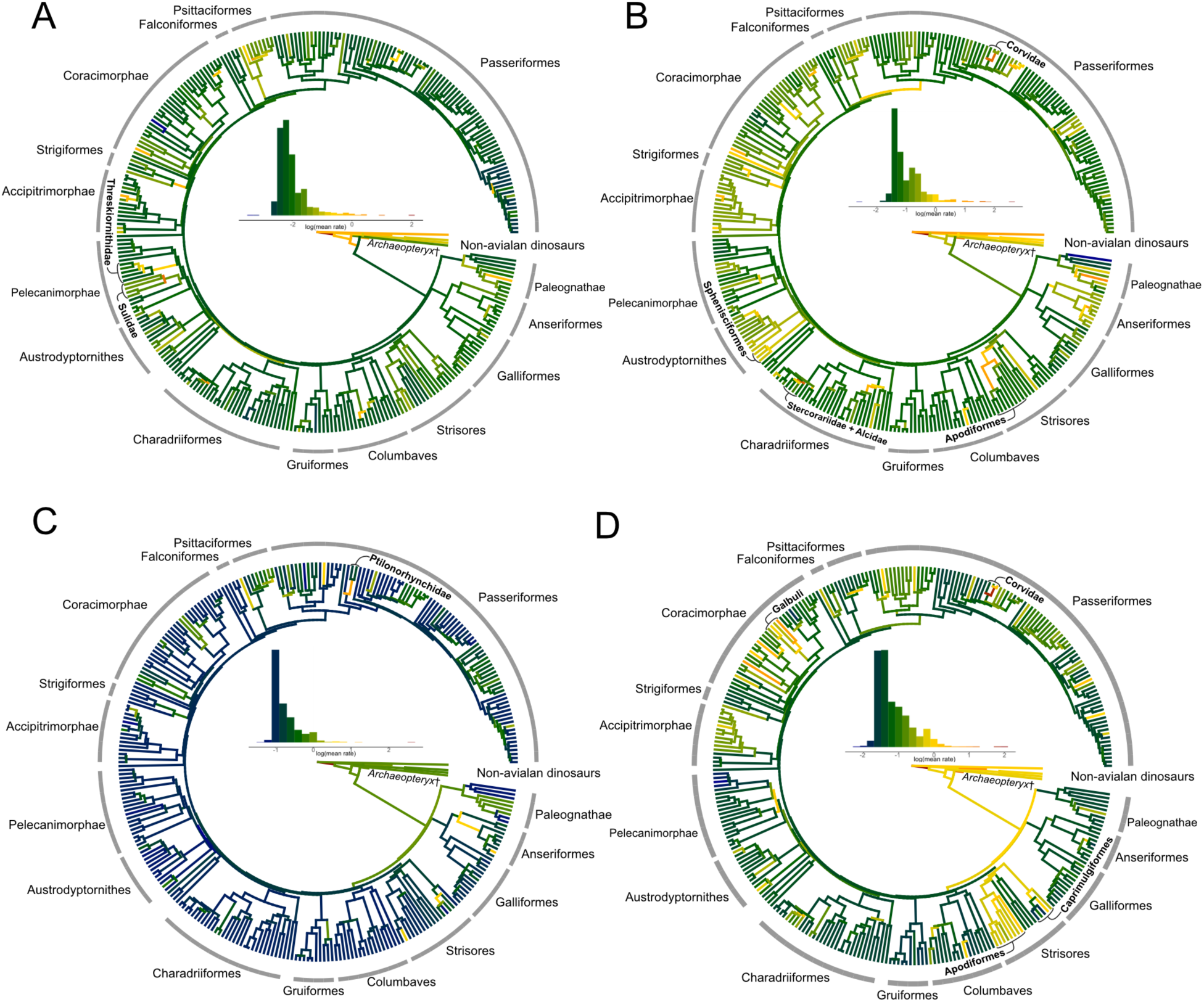
Estimated rates of endocast shape evolution mapped onto time-calibrated phylogeny of birds under best- supported model of trait evolution (λ in all cases), in the cerebrum (A), optic lobe (B), cerebellum (C), and brainstem (D). Color gradient on branches indicates the rate of shape evolution as denoted by the inset histograms. Taxonomic naming from ref. ^146^. Species names on a phylogenetic tree are given in Fig. S3.

### Ecological covariation

Visually, the PC morphospaces and allometry plots (Fig. 1, 2, S1) do not show clear clustering by ecological and developmental traits, except for terrestrial and incessorial foragers (Fig. S1, S2). To test for patterns of ecological covariation with brain morphology, we performed phylogenetically-informed MANOVAs on endocranial size and shape with ecological traits on extant birds (Fig. 4, S5; Table S1). Analyses were conducted on whole endocast shape and its subdivisions (cerebrum, optic lobe, cerebellum, brainstem) against ecological traits (diet, foraging, locomotion, developmental mode) recorded as percentages (diet, foraging) and discrete data (diet, foraging, locomotion, developmental mode) based on previous studies ^40,50^. Neither shape nor size of any endocranial region were found to be significantly correlated with developmental mode when specified as ‘altricial’, ‘precocial’, and ‘semi-precocial’ (*p ≥* 0.2). However, when ‘precocial’ and ‘semi-precocial’ categories were combined into one ‘precocial’ category, brain size relative to body size showed a significant correlation with developmental mode (R^2^ = 0.025, *p* = 0.009), in which altricial birds had larger relative brain sizes than precocial birds in agreement with previous studies^51^. All analyses of shape were corrected for size as part of the multiple linear regression model, which was significantly correlated with shape in each analysis. Each analysis was performed based on Type I sum of squares, in which ecological variables in a model are considered sequentially (e.g., effect of an ecological variable after accounting for evolutionary allometry), as well as on Type II sum of squares, in which all variables were considered together in a single model where one variable is evaluated while controlling for all other variables. This was to differentiate between independent and overlapping effects between ecological variables on morphology, as similar effects on shape between multiple variables can confound explanatory effects in multivariate analyses.

**Fig. 4:**
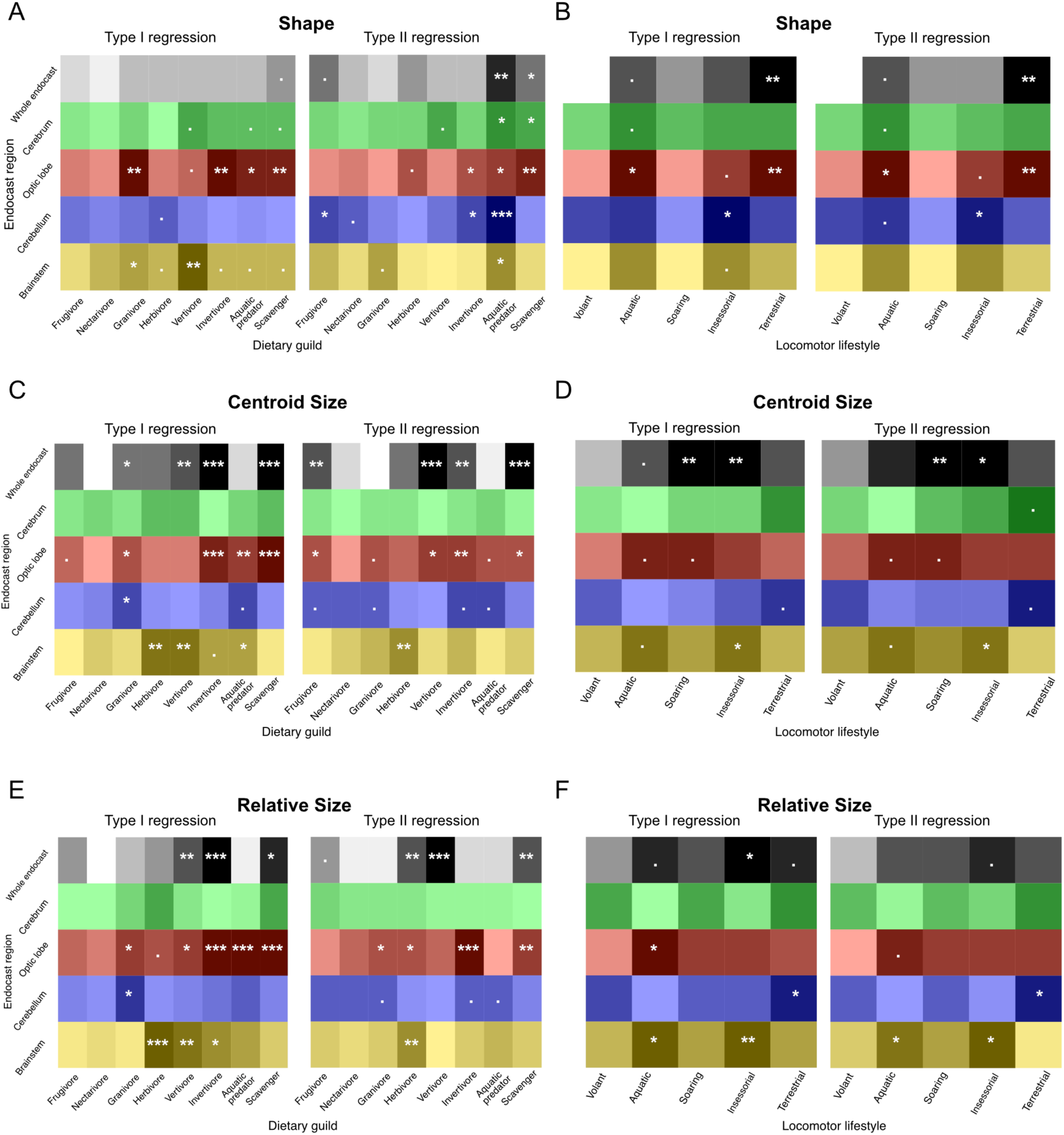
Summary heatmap from ecomorphological analyses on dietary guild and locomotor lifestyle based on Z-scores, where darker gradient indicates greater absolute values, and thus relatively stronger association with endocast shape. Symbols (., *, **, ***) indicate statistical significance based on p-values for each relationship. Gradient is scaled to values along each row. Corresponding values are provided in Table S1.

### Whole Endocast

After correcting for allometry, endocast shape was significantly associated with discrete dietary category (R^2^ = 0.04, Z = 2.14, p = 0.02), as was endocranial size standardized by body mass (R^2^ = 0.097, Z = 3.44, p = 0.002). Across both analyses that were based on Type I sum of squares (only one ecological variable considered at a time after correcting for phylogenetic history and allometry) and Type II sum of squares (all variables considered together and non-sequentially), whole endocast shape was most strongly associated with terrestrial locomotion (Fig. 4; Table S1), in which brains of terrestrial birds are dorsoventrally flatter. Scavenging, which is associated with a flatter dorsal surface of the cerebrum than hindbrain, was more significant in Type II than in Type I analyses, indicating a potential interaction between variables in the model. Aquatic predation was also significantly associated with endocranial shape in Type II but not Type I analyses. Endocast centroid size, used as a metric for absolute brain size, was significantly associated with vertivory, invertivory, scavenging, soaring, and insessorial lifestyle across both Type I and Type II analyses. Granivory was significantly associated with endocast centroid size in Type I but not Type II analyses, suggesting an equivocal association when other variables are accounted for, while frugivory was significant in Type II but not Type I analyses. Endocast size corrected by body mass, used as proxy for brain size relative to body size, was significantly associated with vertivory and scavenging across both Type I and II analyses. Invertivory and aroboreality were significant in Type I analyses but not Type II, while herbivory was significant in Type II but not Type I analyses (Fig. 4; Table S1). Results based on foraging guild were generally compatible with those of diet and locomotion, and thus are described in the Supplementary Text.

### Cerebrum

Individual brain regions were compared with ecological variables to assess regionalized ecomorphological signal. The cerebrum is implicated in many functions, including sensory innervation, integration, and voluntary motor control^23,52–54^. One of three visual pathways in the avian brain, the thalamofugal pathway, terminates in the cerebrum as a protuberance called the ‘Wulst’ on the dorsal surface of each cerebral hemisphere, with various associated functions including binocular vision, motion perception, and spatial orientation^23,25,55–57^. Cerebrum shape was significantly associated with discrete dietary category (R^2^ = 0.034, Z = 1.79, p = 0.036), while cerebrum size was not. Vertivory and aquatic locomotion were weakly associated with cerebrum shape across both Type I and II analyses, while aquatic predation and scavenging were significantly associated with cerebrum shape in Type II analyses but not Type I. Vertivory was associated with the posterior edge of the cerebrum being flatter on the dorsal surface, while the anterodorsal edge, where the Wulst sits, is more bulbous, suggesting an association with stronger binocular vision in vertivores. Aquatic predation was associated with a flatter cerebrum, while scavenging was associated with a rounder cerebrum. Neither cerebrum centroid size nor proportional size, calculated as the size of the cerebrum divided by the total endocast size, were significantly associated with any dietary guild or locomotor variables (Fig. 4; Table S1).

### Optic lobe

The optic lobe is a midbrain structure related to processing visual signal from the retina, and is a major component of the primary visual system, the tectofugal pathway^23,58^. Optic lobe shape was significantly associated with discrete dietary category (R^2^ = 0.045, Z = 2.68, p = 0.002), as was absolute size (R^2^ = 0.22, Z = 3.32, p = 0.002) and proportional size (R^2^ = 0.12, Z= 3.9, p = 0.001). Across both Type I and Type II analyses, optic lobe shape was significantly associated with invertivory, aquatic predation, scavenging, aquatic locomotion, and terrestriality, and weakly with insessorial lifestyle. Granivory was significant in Type I but not Type II analyses (Fig. 4; Table S1). Of these variables, the strongest effects were observed in relation to scavenging, in which optic lobes are flatter and narrower, and in invertivory, in which optic lobes are dorsally expanded and bulbous. Optic lobe centroid size was significantly associated with granivory, invertivory, aquatic predation, scavenging, and weakly with aquatic locomotion and soaring across both Type I and II analyses. Frugivory and vertivory were significant in Type II but not Type I analyses (Fig. 4; Table S1). Proportional optic lobe size was significantly associated with granivory, invertivory, scavenging, and aquatic locomotion across both Type I and Type II analyses. Vertivory and aquatic predation were significant in Type I but not Type II analyses, while herbivory was significant in Type II but not Type I analyses (Fig. 4; Table S1). Among brain regions, optic lobe shape showed the greatest degree of ecomorphological signal based on abundance of significant correlations with ecological variables and their corresponding R^2^ values.

### Cerebellum

The cerebellum, among performing other functions, processes input from both the visual and vestibular systems to control motor functions associated with balance and initiation of muscular movement^59,60^. Despite occupying a small proportion of the avian brain mass, it contains nearly half of all neurons in the brain, and is characterized by substantial infolding^61–63^. Cerebellum shape was weakly associated with discrete dietary category (R^2^ = 0.04, Z = 1.59, p = 0.06), while cerebellum size was not. Across both Type I and Type II analyses, cerebellum shape was significantly associated with insessorial lifestyle, in which the cerebellum is slightly shortened anteroposteriorly but rounded dorsally. Frugivory, invertivory, and aquatic predation were significantly associated with shape in Type II but not Type I analyses, indicating a potential interaction between variables, which is corroborated by the significant association between cerebellum shape and foraging for aquatic invertebrates (Table S2; Fig. S4). Of these, the strongest effects were observed in relation to aquatic predation, in which the cerebellum is anteroposteriorly lengthened. Cerebellum centroid size was weakly correlated with granivory, aquatic predation, and terrestriality across both types of analyses. Invertivory was weakly significant in Type II analyses but not Type I (Fig. 4; Table S1). Proportional cerebellum size was significantly associated with terrestriality, as well as weakly with granivory across both types of analyses. Aquatic predation and invertivory were weakly associated with proportional cerebellum size in Type II but not Type I analyses (Fig. 4; Table S1).

### Brainstem

The brainstem connects the brain to the spinal cord, and is less intrinsically involved in cognitive functions but critical for key involuntary functions (respiration, heart rate) ^60,64,65^. Here, we capture the shape of the hindbrain portion of the brainstem (‘brainstem’ hereafter). Brainstem centroid size (R^2^ = 0.06, Z = 1.72, p = 0.05) and proportional size (R^2^ = 0.07, Z = 2.12, p = 0.02) were significantly associated with discrete dietary category while brainstem shape was not. Brainstem shape was significantly associated with granivory, and weakly with aquatic predation, across both Type I and II analyses (Fig. 4; Table S1). Vertivory was significant, while herbivory, scavenging, and insessorial lifestyle were weakly significant, in Type I but not Type II analyses. Granivory was associated with a brainstem that is mediolaterally narrow and anteroposteriorly elongate, suggestive of a posteriorly positioned foramen magnum, while insessoriality was associated with a shortened and anteroposteriorly arched brainstem, suggesting a more ventral foramen magnum and upright posture. Brainstem centroid size was significantly associated with herbivory, insessorial lifestyle, and weakly with aquatic locomotion across both Type I and II analyses (Fig. 4; Table S1). Vertivory and aquatic predation were significant in Type I but not Type II analyses. Proportional brainstem size was significantly associated with herbivory, aquatic locomotion, and insessorial lifestyle across both Type I and II analyses. Vertivory and invertivory were significant in Type I but not Type II analyses (Fig. 4; Table S1).

### Discriminant analysis

To assess the predictive power of endocranial shape for ecology, we used phylogenetic functional discriminant analysis (pFDA) and performed cross-validation to measure accuracy in assigning ecological traits to extant species. The analysis resulted in70% misclassification for dietary category, suggesting that despite a significant correlation, endocranial shape is a poor predictor of dietary category. All three locomotor variables showed approximately 50% misclassification error. Over 100 replicates of the discriminant function analysis with equivalent within-group sample size, aquatic lifestyle showed the lowest misclassification rate (mean: 43%) and the greatest variance in predicted error (25-53%), suggesting the fewest misclassifications across all replicates. Misclassification was similar for terrestriality (47% [range: 43% - 50%]) and insessorial lifestyle (48% [range: 44% - 50%]). As such, endocast shape has poor predictive power for ecological traits in crown birds.

## DISCUSSION

### Brain Shape Variation and Evolution

The biological significance of avian brain morphology is supported by a long history of comparative studies examining cognitive, behavioral, ecological, allometric, evolutionary, and functional correlates of neuroanatomical variation^5,25,43,66–70^. Much of this work has focused on volumetric variation among brain regions^5,22,71^ or on internal morphological features such as cerebellar foliation and infolding^62,63^, relating such variation to allometry, phylogenetic history, and behavioral specialization. Collectively, these studies demonstrate that morphological variation among avian brains reflects biologically meaningful differences in underlying neuroanatomical organization. However, gross brain morphology is influenced by several interacting factors, including neural architecture, external and internal cranial morphology, sensory structures, and developmental integration^43,59,72,73^. While previous research has largely emphasized volumetric and internal neuroanatomical variation, our results suggest that three-dimensional brain shape represents an additional and previously underexplored axis of morphological variation that exhibits ecologically meaningful variation across birds.

The elongate-globular spectrum drives the variation in avian brain shape. This result is consistent with the spatial packing hypothesis of brain evolution^35^, in which reorganization of brain region configuration is associated with changes in cranial flexion and foramen magnum position, and is largely independent of variation in size metrics (Fig. 2). This flexion of the cranial base and the reorganization of the brain into a more globular structure is accompanied by a transition to a more ventrally oriented foramen magnum observed across both birds and mammals^34,35,37,39,74,75^. Conversely, the elongated brain morphotypes are characterized by a smaller and more anteriorly positioned cerebrum with reduced dorsal curvature, a feature convergent between aquatic birds and non-avian dinosaurs, which possess a posteriorly positioned foramen magnum. Since foramen magnum position directly influences head posture, our results imply that head posture differences are also likely to be strongly associated with brain shape variation in crown birds. This outcome agrees with a study by Kawabe and colleagues^38^, which found that birds with a posteriorly oriented foramen magnum also possess more anteroposteriorly elongated brains accompanied by elongated and laterally-oriented orbits, whereas birds with globular brains and orbits have a ventrally positioned foramen magnum and a more upright posture. Likewise, Kulemeyer and colleagues^76^ suggest that birds with a more posteriorly oriented foramen magnum and anteroposteriorly elongated cranium (and thereby endocast) experience reduced drag during sustained locomotion. A posterior foramen magnum implies a longer neck to facilitate hydrodynamic streamlining and to hold the head upright; neck length has been shown to scale isometrically with both body mass and head mass^77^, which agrees with our results that show this brain morphotype is more common in larger birds. Our shape analysis shows that these globular and elongated morphotypes are partly associated with aerial and aquatic foragers, respectively, which have adapted for an upright, perching stance in the former and a hydrodynamic body profile in the latter.

Evolutionary rates analyses reveal accelerated evolution among non-avian coelurosaurian dinosaurs and highly heterogeneous trends across avian taxa and among brain regions, complementing previous macroevolutionary studies on avian brain size evolution (e.g., ^5^). While this rates analysis differs from identifying distinct shifts in brain size parameters, elevated rates of brain shape evolution mirrors many of the phylogenetic positions where distinct allometric trends were identified previously ^5^. These include the lineage leading to non-avian dinosaurs, *Apteryx* (kiwi), Apodiformes (hummingbirds and swifts), Strigiformes (owls), Picidae (woodpeckers), Psittaciformes (parrots), Ptilonorhychidae (bowerbirds), and Corvidae (crows, ravens). Especially noteworthy are the elevated rates of morphological evolution in two exemplary passerine groups, the bowerbirds and the corvids. Bowerbirds are notorious for their complex nest-building and courtship strategies ^78^ and previous studies found a positive relationship between the complexity of their bowers and brain size, specifically cerebellum size ^79,80^. Our analyses show accelerated evolutionary rates in cerebellum shape in bowerbirds, where the cerebellum is anteroposteriorly shortened, dorsoventrally rounded, and ventrally angled. Together, these results suggest an association between the cognitive traits that are associated with a proportionately larger and arched cerebellum, such as motor planning and control, procedural learning, postural displays in bower construction behaviour, and a greater motor than visual cognitive load^80^. Similarly, cerebrum, optic lobe, and brainstem shapes show elevated evolutionary rates in the lineage leading to the highly encephalized Corvidae^81^. The brains of corvids have been extensively investigated in relation to their cognitive abilities, with several studies on their relatively large brains^12,82^, and specifically on the size of associative and motor-related structures in the forebrain associated with tool use and problem-solving skills in New Caledonian crows^11,83–85^. The rapid evolution towards a shorter brainstem in corvids could be explained by a number of factors, including a more ventral foramen magnum position and thus postural stance that facilitates controlled motor behaviors^76^, or spatial packing of a large cerebrum leading to restructuring of the remaining brain regions^39^.

### Ecomorphological Trends in Avian Brain Shape Evolution

Taxa exhibiting the two extremes of the elongated-globular spectrum also show some of the strongest associations with ecology. Aerial foraging specialists like Strisores, including Apodiformes (hummingbirds and swifts) and Caprimulgiformes (nightjars and nighthawks) show the most globular and anteroposteriorly shortened forebrains, and ventrally angled hindbrains. These groups are uniquely adept at intricate maneuvers in flight to either catch insect prey^86^ or hover over nectar-bearing flowers^87,88^, and further display elevated rates of shape evolution in the brainstem^65^. Caprimulgiforms also show distinct convergent traits with owls, notably their wide binocular visual fields along with extremely large Wulsts^28,89,90^, however owls do not show nearly as globular brains as caprimulgiforms do. This suggests that patterns of globularization are neither driven by enlarged Wulsts nor relative brain size among birds more broadly. The other end of the spectrum is occupied by the aquatic predatory Pelecaniformes, particularly the anhinga and cormorant, which showed anteroposteriorly elongated and straight endocasts. All aquatic predators have this brain morphotype to some extent, likely due to the necessity for a hydrodynamic body profile, including of the skull, in an aquatic predatory lifestyle^40,91–94^. As such, the elongated brain of these taxa may be driven by adaptations in the skull for an aquatic lifestyle, suggesting a high degree of integration between the brain and skull.

Several bursts of brain shape evolution co-occur with major ecological transitions. While both MANOVA and pFDA indicate that avian-wide associations between ecology and brain shape are weak overall, analysis of evolutionary rates points to clade-specific evolutionary patterns that could be strongly linked to their specific ecologies. For instance, the lineage leading to Strisores (‘nightbirds’) exhibit fast evolution in the optic lobe which could reflect adaptation for maintaining visual acuity for nocturnal foraging. Within Strisores, Apodiformes (hummingbirds, swifts), which engage in more diurnal habits, show subsequent accelerated rates of evolution in the optic lobe. Indeed, the endocasts of most non-apodiform Strisores that we sampled show more quadrangular optic lobes that extend more posteriorly along the ventral margin of the cerebrum, whereas apodiform taxa possess optic lobes that are more spherical and positioned anteroventrally to the cerebrum, as is more typical in birds. Rapid evolutionary changes in the optic lobe also occurred towards the nocturnal *Apteryx* (kiwi), but in contrast to the nocturnal Strisores species (e.g., nightjars), their optic lobes are highly reduced in favor of reliance on their olfactory sense^95^. Similarly, the optic lobes underwent rapid evolution independently among raptors which rely on vision for their predatory behavior. Notably, the cerebrum shape also underwent rapid changes independently in the polyphyletic Old and New World vultures, which are likely attributed to enlarged olfactory bulbs that are continuous with the anterior portion of the cerebrum. Accelerated changes to the cerebrum are observed for Threskiornithidae (spoonbills, ibises) with hypersensitive beaks, that could be associated with increased sensory perception through the beak in this clade. Their cerebra show a hypertrophied anterior portion near the olfactory bulbs that receives somatosensory signals, including those from trigeminal nerves that provide sensory innervation to the beak^96,97^. This result is consistent with larger principal sensory nucleus of the trigeminal nerve in the brainstem observed in birds that rely on specialized sensory abilities using their beak or tongue^73^. The cerebellum underwent moderately fast shape evolution in the lineage leading to Rallidae (rails), a clade with multiple secondary losses of flight. However, the shape changes are unlikely to be due to flightlessness as the sampled rallid species are volant, and previous studies have shown poor correlation between brain morphology and flightlessness^48,98,99^. Interestingly, omnivory showed high rates of evolution in cerebellum and second highest in cerebrum which is consistent with previous studies that have shown correlation between larger brains with more generalist diets^100^.

Developmental mode, defined by the distinction between altricial and precocial development, was significantly correlated with relative brain size, with altricial birds possessing larger relative brain sizes. Substantial evidence has supported the hypothesis that altricial birds tend to have larger brains, as a longer growth period and greater parental investment allows for further brain development after hatching^22,51,101,102^. The lack of correlation between developmental mode and brain shape, independent of size, is validated by previous findings that skull shape is also not significantly correlated with developmental mode^50^. This suggests that while a longer post-hatching growth period allows for greater brain volumes to develop, brain shape is not uniformly associated with timing of ontogenetic development across birds.

### Similarities and Differences in Brain Shape and Size Evolution

Endocast size, and by extension brain size, accounts for a very small percentage (∼6%) of brain shape variation after correcting for phylogenetic structure. Therefore, the majority of the variation in brain morphology is unaccounted for when size or volumetric data are considered alone. Ksepka et al.^5^ identified parrots and crows as having independently enlarged relative brain sizes, followed closely by owls, woodpeckers, and bowerbirds. Our analyses of shape evolution revealed elevated rates in some of these groups for certain brain regions, including the whole endocast and cerebrum shapes in parrots, which possess an inflated cerebrum with enlarged Wulsts, an anteroposteriorly shortened and rounded cerebellum in bowerbirds, and anteroposteriorly shortened and dorsoventrally flattened brainstem in crows. Groups with substantial evolutionary shifts in brain shape that did not correspond with associated size evolution included the two extremes of the elongated-globular spectrum: the overall highly globular brain and laterally rounded optic lobe of the Apodiformes (hummingbirds and swifts), and the elongated brains with flatter optic lobes in the aquatic predators in Pelecaniformes. Interestingly, optic lobe shape evolution, which showed the highest correlation with ecology, did not result in evolutionary shifts that corresponded with brain size enlargement. This reinforces the ecological significance not only of the elongated-globular spectrum, but of the optic lobe itself, which is rounder and dorsoventrally elongated in more aerial birds (e.g, those with insessorial and aerial foraging ecologies), while being flatter (less convex) and laterally oriented in terrestrial and aquatic birds. The optic lobe is also enlarged in long-distance migratory birds^103^, while being proportionately smaller in waterfowl, owls, and parrots despite no evidence of reduced visual ability through the tectofugal pathway^68^. Between the two visual pathways that occupy space on the brain surface, the optic lobe shape appears to show significantly more functional and ecological signal than the Wulst. Therefore, while relative size loosely correlates with this elongated-globular spectrum of shape (Fig. 2), crows and parrots – which have been touted as the birds with the largest relative brain sizes and greatest cognitive ability – do not fall under either extreme of elongated or globular brains. For example, parrot brains have distinctly quadrangular brain shapes and a pronounced interhemispheric sulcus in larger parrot taxa^104^, suggested to accommodate a bony interhemispheric septum between their olfactory bulbs, which may provide protection to the brain during feeding and locomotion^105^. However, parrots did not display any consistent shape traits associated with ecology, and their large and specialized beaks are more strongly associated with allometry and integration rather than with diet^106^. However, parrots do possess enlarged Wulsts, which in conjunction with the optic lobe, has been associated with processing brightness and visual pattern discrimination^107,108^. Overall, while several ecomorphological trends in brain shape and size correspond with one another, brains that show extreme differences in elongated or globular shapes possess ecological and evolutionary patterns that are more independent of size variation, demonstrating that shape variation is a significant component of neuroanatomy that is subject to selective forces.

While there is significant overlap in brain shape across clades, our complementary analyses of endocast size and shape reveal large-scale patterns in avian brain evolution: 1) after accounting for evolutionary allometry, species with primarily aerial-based ecologies have more globular brain shapes, while terrestrial and aquatic birds have more elongated brains; 2) optic lobe shape has the greatest number of ecological covariates and the greatest variation in evolutionary rates; and 3) despite statistically significant correlations, predictive power of brain shape for ecology is limited when applied across birds. The poorly predicted form-and-function link does not demonstrate the absence of a strong connection between brain morphology and functional abilities—rather, that the relationship is complex across the entire avian phylogeny. The weak association across macroevolutionary timescales could be due to many-to-one or one-to-many relationships between neuroanatomy, particularly surficial topology, and their ecologically relevant functions^109^. For example, the optic lobe had the greatest number of covariates among brain regions, possibly because the optic lobe has a more targeted function than the other regions characterized in this study. In contrast, cerebrum shape which includes the Wulst, an important structure for visual cues^110^, and involves many other ecologically relevant functions, shows the weakest relationship with ecological variables despite being one of the major drivers of differentiation in brain shape between avian groups. As such, inferring the cognitive and functional ability from endocasts of extinct taxa, for which we have limited behavioral, ecological, and life history information should be approached with caution. Overall, we find evidence of foramen magnum position and hydrodynamic streamlining having strong influence on brain shape, which complements the findings of previous studies on integration between endocranial shape and surrounding skull structures^38,111,112^. Using comprehensive data on brain morphology, our analyses show that the remarkable neuroanatomical diversity in modern birds emerged not through the dominance of any particular factors, but rather through independent divergence of brain structures and avian clades from the global allometric and ecomorphological patterns.

## METHODS

### Endocranial Sampling

We compiled endocranial reconstructions for 300 extant avian species that encompass their phylogenetic, ecological, and morphological breadth (Supplementary Text). Of the 300 extant samples, 124 endocast models were created from CT imaging of avian specimens by the authors (mostly skeletal, but fluid specimens were used when skeletal were not available), 96 endocasts models were newly constructed from CT image stacks or mesh files available on Morphosource ^113^, and 80 endocast mesh files were created and shared by colleagues. For anchoring the ancestral condition, we also included endocasts from six non-avian coelurosaurian dinosaur fossils from previous studies^114,115^: *Tyrannosaurus rex* (FMNH PR2081)*, Incisivosaurus gauthieri* (IVPP V 13326)*, Citipati osmolskae* (IGM 100/973)*, Khaan mckennai* (IGM 100/973), an unnamed troodontid IGM 100/1126, and *Archaeopteryx lithographica* (NHMUK PV OR 37001). Multiple methods were used for segmentation of endocranial cavity (see Supplementary Text for additional information). Due to the broad phylogenetic scope of this study, any systemic differences in shape data due to different imaging, segmentation, and mesh creation procedures were considered to be negligible compared to interspecific variation. We refer to endocranial regions by the name of the brain structure that left the impression (e.g., ‘cerebrum’ for the impression of cerebrum on the endocast).

### Morphometric data

To quantify shape of the endocast, we collected high-density GM data building off the landmark scheme of ^44^ (Fig. S5; Table S3). We used Stratovan Checkpoint ^116^ to digitally place landmarks and curve semilandmarks on the right side of the endocasts, capturing the major brain regions, including the cerebrum, optic lobe, cerebellum, and brainstem. Additional sliding semilandmarks on the surfaces of these brain regions were semiautomatically placed by implementing the ‘placePatch’ function from the R package ‘Morpho’ version 2.12 ^117^. This function uses a template, in this case from the model avian species *Gallus gallus*, consisting of an endocast mesh and corresponding landmark and semilandmark curve coordinates, along with a constellation of evenly distributed surface semilandmarks, and maps these surface semilandmarks for each endocast mesh. To limit biases produced by performing Procrustes alignment on one-sided data of bilaterally symmetrical structures ^118,119^, the right sided landmark configuration was mirrored along the median plane to produce a bilaterally symmetrical dataset of endocast landmarks using the ‘mirrorfill’ function in the ‘paleomorph’ R package ^120^. Using the ‘slider3d’ function in the package ‘Morpho’^117^, all curve and surface semi-landmarks were then slid along their corresponding mesh surface, while minimizing total bending energy ^121,122^. Generalized Procrustes alignment (GPA) was performed using the ‘gpagen’ function in the ‘geomorph’ R package to generate Procrustes shape coordinates and extract centroid sizes ^123^. Aligned coordinates from the mirrored left side of the brain were then removed, and subsequent analyses were conducted only on coordinates from the midline and right side of the brain to avoid overweighting of bilateral landmarks through duplicated shape information. This procedure resulted in a total of 241 landmarks, of which there were 13 fixed landmarks, 130 curve semilandmarks and 98 surface semilandmarks. For regional analyses, we extracted a subset of globally aligned Procrustes coordinates that characterize each brain region and ran another round of GPA to locally align the coordinates of each region. In addition to centroid size, proportional regional size of each brain region was calculated as the centroid size of the region in question divided by the total endocast centroid size. Body mass data were taken from ^124^.

### Ecological data

To test ecomorphological hypotheses, we compiled a set of ecological traits from previously published categorizations and ecological assessments, primarily drawing from the global standard dataset AVONET^40,125–127^ (Table S4). These were designated as: 1) discrete dietary categories, 2) dietary guild scores, 3) foraging guild scores, 4) locomotor traits, and 5) developmental modes. Discrete dietary categories (frugivore, nectarivore, granivore, herbivore, vertivore, invertivore, aquatic predator, scavenger) were assigned based on whether each taxon obtained greater than or equal to 60% of its resources from any one food category, with remaining taxa being classified as ‘omnivores’, in which no single food category constituted ≥60% of total intake (Table S4). Scavengers are distinguished from land-based predatory vertivores to account for the distinct neurosensory adaptations related to the detection and consumption of carrion and animal carcasses^128–130^. These are coarse categorizations and were primarily used for visualization purposes in figures 1 and 2, as well as in the Discriminant Analysis (see below). Dietary niche scores (which match the discrete categories except for ‘omnivore’; Table S1, S4) and foraging guild scores (Table S2) were classified as continuous percentages of total dietary intake (0-100%), to broadly characterize the trophic niche of each taxon. These foraging guilds encompass both dietary resource and the method by which the species obtains the resource and are thus not independent from the dietary scores, but broadly capture additional traits to encompass ecological complexity. These percentage data were tested in Type I PGLS analyses (see ‘Ecomorphological Analysis’ below), and were then converted to binary presence/absence data for multivariate (Type II) PGLS analyses, to avoid variable redundancy. Locomotor traits were characterized to encompass the full spectrum of avian lifestyle ecology, scored as binary data (0/1) for the following traits: volant, aquatic lifestyle, soaring, terrestrial, and insessorial^131^ (Table S4). Finer sub-classifications of powered flight (e.g., intermittent bounding or flap-gliding) were omitted to maintain model parsimony, as these styles share overlapping morphological profiles and lack distinct anatomical or sensory signatures in macroevolutionary comparative analyses (e.g., ^132,133^). Finally, categorical data on developmental modes (altriciality, semi-precociality, precociality) were compiled from ^50^ to examine the relationship between brain shape and life history strategies.

### Phylogeny

Phylogenetic comparative analyses were conducted using a composite extant phylogeny from ^134^, which combines the genus-level resolution from the supertree distribution of the ^135^ backbone, and scales the age estimates and branch lengths according to ^136^. We incorporated the fossil taxa to the base of the extant tree using an equal branching method ^1^ based on the middle age of the occurrence estimates from the Paleobiology Database (paleobiodb.org).

### Analysis

#### Morphometric variation

We conducted comparative morphometric analyses using the R package ‘geomorph’ ^123^ unless indicated otherwise, performed in R version 4.3.2 ^137^. To visualize patterns of morphological variation in the avian brain of the extant + fossil dataset, major axes of shape variation were visualized using a principal component analysis (PCA) on the Procrustes coordinates, using the ‘gm.prcomp’ function. Phylogeny was mapped onto the PCA to create a ‘phylomorphospace’, to aid in visualization of the phylogenetic structure in shape variation. We tested for phylogenetic signal of endocranial shape for the entire endocast, as well as for each region, using Blomberg’s K test-statistic through the ‘physignal’ function^138^.

We evaluated allometric patterns of variation for both endocranial shape and size in the extant dataset alone, as well as in the extant + fossil dataset, with centroid size of the endocast as a proxy for brain size ^139^. Allometry of brain size was analyzed by comparing endocast centroid size with body mass from ^124^ of each taxon from the extant dataset, analyzed with a phylogenetic generalized least squares (PGLS) regression using the ‘gls’ function from the R package ‘nlme’ version 3.1-166 ^140^. We analyzed allometry of endocast shape by performing a phylogenetic generalized least squares (pGLS) analysis on three variable combinations: 1) shape against log centroid size of the extant-only dataset, 2) shape against log-cube rooted body mass of the extant-only dataset, and 3) shape against log centroid size of the extant + fossil dataset. These allometric relationships were visualized by plotting the given size variable against regression score coefficients, which are based on an eigenvector that maximally varies accordingly with the independent variable in question (e.g., the size variable), accounting for the maximum correlation between shape changes predicted by the regression model^141,142^. Since endocast shape was significantly correlated with endocast size (see results), centroid size was incorporated in linear models used in subsequent ecomorphological analyses.

### Rates of evolution

To estimate evolutionary rates for the overall brain and its partitions, we used BayesTraits V4 ^49^ to fit variable-rates evolutionary models to the extant + fossil dataset. Rates were calculated on PC axes accounting for 99% of the total brain shape variation were analyzed. All available evolutionary models were compared, including BM, OU, δ, κ, and λ models, and the best- supported model was determined using Bayes Factor comparisons using functions from the BTRtools R package (source code archived at github.com/tiago-simoes/BTprocessR2). Rates analyses were performed on the total endocast shape, and then on each individual brain region captured by the endocast (cerebrum, optic lobe, cerebellum, brainstem). Further, we compared relative evolutionary rates within each brain partition to examine how they differ between extant birds with different dietary categories. This was implemented with the ‘compare.evol.rates’ function in ‘geomorph’, and then rates were divided by the highest rate in the brain region to standardize rate values and compare relative rates for each dietary category (Extended Results, Fig. S6).

### Ecomorphological analysis

We tested for ecological correlates of endocast shape of only extant birds using distance-based phylogenetic generalized least squares (PGLS) on the Procrustes coordinates. These were conducted using the ‘procD.pgls’ function, which performs phylogenetically corrected Procrustes distance-based regression analysis under a model of Brownian motion, by analyzing pairwise distances between observations rather than estimating a covariance matrix, which retains high statistical power when the number of variables exceeds the number of observations^143^. We performed two sets of analyses, one using the Type I (sequential) sum of squares for analyses of each ecological variable separately, and Type II (hierarchical) sum of squares on a set of ecological variables^144^. Separate analyses were conducted for each brain region: whole endocast, cerebrum, optic lobe, cerebellum, and brainstem. Shape and size for each region were compared against discrete dietary category, and then further sets of regression models were implemented for each set of multivariate data: 1) dietary guild data, 2) foraging guild data, and 3) locomotor ecology data (Table S1, S2). For dietary guild scores and foraging guild scores, Type I analyses used percentage score data and included one ecological variable per analysis, while Type II analyses incorporated all variables into a single model, and converted these data into presence and absence of each category. If a variable was found to be statistically significant in a Type I analysis but not Type II, then the correlation is considered to be equivocal; alternatively, if the variable is significant in Type II but not Type I, then there is likely an interaction or strong covariation between ecological variables in the model that are associated with the morphological variable when considered together.

### Discriminant analysis

To test for the predictive power of endocranial shape for dietary trait, we implemented a phylogenetic flexible discriminant analysis (pFDA) on the Procrustes coordinates of the extant dataset using the ‘phylo.FDA’ function developed by ^145^. Similar to linear discriminant analysis, pFDA uses categorical data, such as dietary category, and multivariate shape data to estimate the posterior probability of ecological traits for each taxon while accounting for phylogenetic non-independence in the multivariate dataset. Since there is unequal representation of each dietary category in the dataset, we subsampled the dataset at random to reach an equal number of taxa exhibiting each trait before running the pFDA and repeated this for 100 replicates to obtain robust estimates for the strength of affinity for each ecological trait.

