## Supplementary Text for "Complementary ecomorphological patterns in brain shape and size evolution across the avian radiation"

Extended Methods

Fifty of the newly generated CT image stacks were processed by segmenting skulls in Dragonfly and then automating endocasts segmentation with the ‘endomaker’ function in the R package ‘Arothron’ version 2.0.3 ^1,2^, specifying skull mesh decimation to 500,000 triangles, and 200 points of view in endocast construction. While this pipeline produced decent quality endocasts that contain surfaces of regions being landmarked for many specimens, this did not result in full endocranial reconstruction, especially with a gap in the anteroventral surface of the cerebrum between the two laterosphenoid bones in the braincase. Therefore, manual segmentation was performed for newly imaged specimens for full endocranial reconstructions. These consisted of 170 that segmented manually in the software Amira version 2023.2 (ThermoFisher Scientific, Waltham, MA, USA from CT image data. Of the 170 manually segmented specimens, 124 museum specimens were scanned using a Bruker SkyScan 1173 micro-CT scanner at the New York Institute of Technology Visualization Center, and reconstructed using the associated NRecon software to convert scans into cross-sectional images before manual segmentation in Amira.

**Extended Results**

**Evolutionary rates: Relative rates within each region.** We also compared mean evolutionary rates under Brownian motion within each brain partition to examine how they differ between extant birds with different diets (Fig. S6). Carrion scavenging showed the highest rates in the whole brain, cerebrum and optic lobe. Feeding on aquatic animals was associated with highest rates in the brainstem followed closely by vertivory, while omnivory and granivory was associated with the highest rates in the cerebellum. Vertivory and invertivory were generally associated with slower rates across all regions, with the exception of brainstem in vertivorous taxa.

**Ecological covariation: Foraging guild.** For a more detailed characterization of trophic niche, foraging guild categories were scored based on both dietary content and lifestyle/locomotor ecology associated with dietary acquisition. Across both Type I and Type II analyses, overall brain shape was associated with foraging guilds involving invertivory as well as a terrestrial lifestyle, such as foraging for aquatic invertebrates from the ground (Type I: R^2^ = 0.02, Z = 2.94, p = 0.004; Type II: R^2^ = 0.0075, Z = 2.37, p = 0.009), gleaning invertebrates from terrestrial vegetation (Type I: R^2^ = 0.007, Z = 1.92, p = 0.03; Type II = R^2^ = 0.006, Z = 1.52, p = 0.05), and as well as diving for aquatic prey (Type I: R^2^ = 0.008, Z = 2.29, p = 0.013; Type II = R^2^ = 0.006, Z = 1.52, p = 0.05) foraging for fruit on the ground (Type I: R^2^ = 0.0012, Z = 2.83, p = 0.005; Type II: R^2^ = 0.008, Z = 2.34, p = 0.011). Aerial screening for vertebrates (Type I: R^2^ = 0.0013, Z = 2.34, p = 0.009) and sallying to substrate for vertebrates (Type I: R^2^ = 0.006, Z = 1.82, p = 0.0034) was associated with shape in Type I analyses but not Type II, suggesting a weaker association. Aerial sallying for invertebrates (Type I: R^2^ = 0.006, Z = 1.74, p = 0.04) and foraging for terrestrial vegetation (Type I: R^2^ = 0.007, Z = 2.24 p = 0.013) were significant in Type II but not Type I analyses, indicating an interaction between variables (Table S2). Endocranial centroid size was also most associated with terrestrial foraging strategies across both Type I and Type II analyses, including ground frugivory, ground scavenging, and gleaning for terrestrial vertebrates. Sallying to substrate for invertebrates was significant across both types as well. Those significantly associated in Type I but not Type II analyses, indicating a weak correlation, include aerial screening as well as aerial sallying for invertebrates, aquatic predation on water’s surface, aerial-to-substrate predation on vertebrates, and gleaning for arboreal vertebrates; foraging for aquatic vegetation was significantly associated with Type II but not Type I analyses, indicating an interaction between variables. Relative endocranial size showed greatest associations across both Type I and Type II analyses for aerial sallying for invertebrates, sallying to substrate for invertebrates, aerial nectarivory, terrestrial scavenging, and gleaning for arboreal vertebrates (Table S2).

Across both types of analyses, cerebrum shape was most strongly associated with foraging for aquatic invertebrates from the ground (Type I: R^2^ = 0.01, Z = 2.52, p = 0.006; Type II: R^2^ = 0.006, Z = 1.83, p = 0.028). Diving for aquatic prey (R^2^ = 0.008, Z = 2.25, p = 0.011), ground frugivory (R^2^ = 0.008, Z = 1.65, p = 0.043), and aerial screening for vertebrates (R^2^ = 0.012, Z = 2.14, p = 0.018) were significant in Type I but not Type II analyses. Cerebrum centroid size was associated with aerial sallying for invertebrates and sallying to substrate for vertebrates in Type I analyses, and with foraging for both terrestrial and aquatic vegetation in Type II analyses. Proportional cerebrum size was significantly associated with foraging for both terrestrial and aquatic vegetation and gleaning for arboreal vertebrates across both Type I and Type II analyses, while also being associated with ground frugivory and aerial screening for vertebrates in Type II analyses.

Optic lobe shape was associated with foraging for aquatic invertebrates from the ground (Type I: R^2^ = 0.01, Z = 2.09, p = 0.022; Type II: R^2^ = 0.006, Z = 1.69, p = 0.046), aquatic predation from the ground (Type I: R^2^ = 0.02, Z = 2.54, p = 0.01; Type II: R^2^ = 0.008, Z = 2.21, p = 0.013), diving for aquatic prey (Type I: R^2^ = 0.01, Z = 2.82, p = 0.003; Type II: R^2^ = 0.009, Z = 2.36, p = 0.008), foraging for terrestrial vegetation (Type I: R^2^ = 0.007, Z = 1.95, p = 0.023; Type II: R^2^ = 0.008, Z = 2.17, p = 0.02) across both Type I and Type II analyses. Aerial screening for invertebrates, aquatic predation from the air, nectar gleaning, terrestrial granivory, aerial screening for vertebrates, aerial-to-substrate predation on vertebrates, sally-to-substrate predation on vertebrates and ground scavenging were all significant in Type I analyses but not Type II; aquatic scavenging was significant in Type II analyses but not Type I. Across both types of analyses, optic lobe centroid size was most significantly associated with terrestrial gleaning for invertebrates, ground frugivory, foraging for aquatic vegetation, and ground scavenging, while optic lobe proportional size was most significantly associated with gleaning for terrestrial invertebrates, foraging for aquatic vegetation, sally-to-substrate vertebrate predation, and terrestrial scavenging (Table S2).

Cerebellum shape was most significantly associated with foraging for aquatic invertebrates from the ground (Type I: R^2^ = 0.05, Z = 3.39, p = 0.001; Type II: R^2^ = 0.015, Z = 3.18, p = 0.001) and ground frugivory (Type I: R^2^ = 0.027, Z = 3.22, p = 0.001; Type II: R^2^ = 0.015, Z = 2.92, p = 0.002) across both types of analyses. Aerial screening for vertebrates (R^2^ = 0.011, Z = 1.87, p = 0.03) was associated with cerebellum shape in Type I but not Type II analyses; aerial sallying for invertebrates, (R^2^ = 0.014, Z = 2.96, p = 0.003), gleaning for terrestrial invertebrates (R^2^ = 0.009, Z = 2.33, p = 0.012), diving for aquatic prey (R^2^ = 0.016, Z = 2.72, p = 0.001), and gleaning fruit (R^2^ = 0.007, Z = 1.83, p = 0.038) were significantly associated with Type II but not Type I analyses. Cerebellum centroid size was most significantly associated with gleaning for terrestrial invertebrates, feeding on aquatic invertebrates from the ground, aerial screening for vertebrates, and sally-to-substrate vertebrate predation across both types of analyses. Aquatic predation from the air and ground frugivory were associated with cerebellum centroid size in Type I but not Type II analyses. Proportional cerebellum size was significantly associated with gleaning for terrestrial invertebrates and feeding on aquatic invertebrates from the ground across both types of analyses. Aquatic predation from the air was significant in Type I but not Type II analyses, while aerial screening for vertebrates was significant in Type II but not Type I analyses.

Brainstem shape was most significantly associated with foraging for aquatic invertebrates from the ground (Type I: R^2^ = 0.016, Z = 2.3, p = 0.014; Type II: R^2^ = 0.008, Z = 2.03, p = 0.023), diving for aquatic prey (Type I: R^2^ = 0.008, Z = 1.83, p = 0.032; Type II: R^2^ = 0.015, Z = 3.14, p = 0.001), aerial-to-substrate predation on vertebrates (Type I: R^2^ = 0.044, Z = 3.47, p = 0.001; Type II: R^2^ = 0.016, Z = 3.32, p = 0.001), and sally-to-substrate predation on vertebrates (Type I: R^2^ = 0.02, Z = 2.99, p = 0.004; Type II: R^2^ = 0.01, Z = 2.6, p = 0.008) across both types of analyses. Gleaning for arboreal invertebrates (R^2^ = 0.01, Z = 2.52, p = 0.007), aquatic predation from the ground (R^2^ = 0.016, Z = 2.3, p = 0.014), aquatic predation from the air (R^2^ = 0.027, Z = 2.94, p = 0.002) and aerial screening for vertebrates (R^2^ = 0.069, Z = 3.16, p = 0.001) were associated with shape in Type I but not Type II analyses; aerial screening for invertebrates (R^2^ = 0.009, Z = 2.39, p = 0.008), aerial sallying for invertebrates (R^2^ = 0.01, Z = 2.57, p = 0.002), sallying to substrate for invertebrates (R^2^ = 0.006, Z = 1.7, p = 0.047) were significant in Type II but not Type I analyses. Brainstem centroid size was significantly associated with sally-to-substrate predation on vertebrates across both types of analyses. Aerial screening for invertebrates, aquatic predation from the ground and the air, foraging for aquatic vegetation, and aerial screening as well as aerial-to-substrate predation on vertebrates were significant in Type I but not Type II analyses; aerial sallying for invertebrates, gleaning for fruit and terrestrial granivory were significant in Type II but not Type I analyses. Proportional brainstem size was significantly associated with ground frugivory and sally-to-substrate predation on vertebrates across both types of analyses. Aerial screening for invertebrates, aquatic predation from the air, foraging for aquatic vegetation, and aerial screening and aerial-to-substrate predation on vertebrates were significant in Type I but not Type II analyses; gleaning fruit, terrestrial granivory, and feeding on terrestrial vegetation were significant in Type II but not Type I analyses (Table S2).

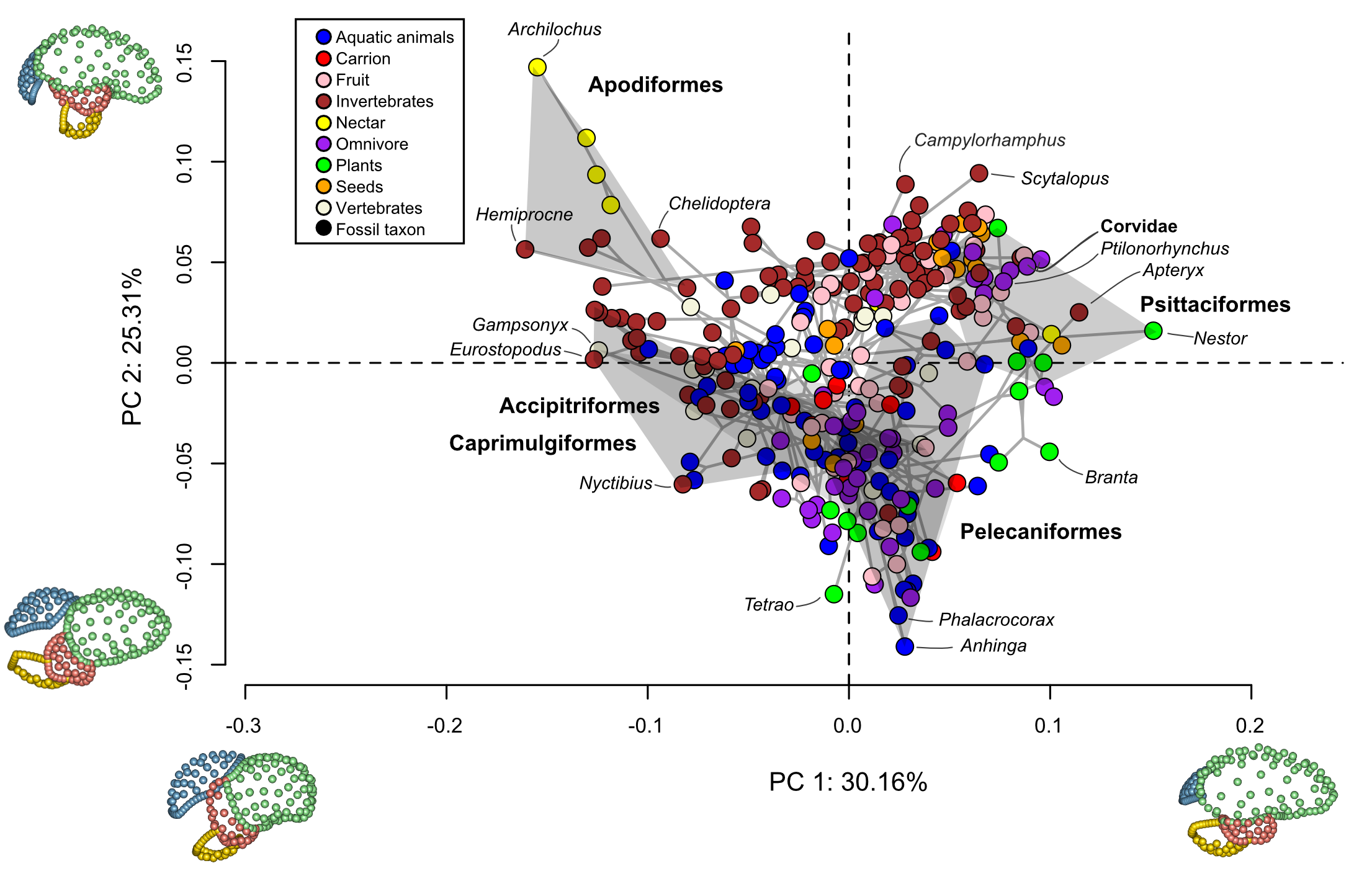

**Fig. S1:** Phylomorphospace of the first two principal component (PC) axes highlighting the major aspects of endocranial shape variation in extant taxa only. Points are colored by discrete dietary category. Inset images along the axes depict shape changes associated with PC1 and PC2 axes, colored by region (green; cerebrum; red: optic lobe; blue: cerebellum; yellow: brainstem)

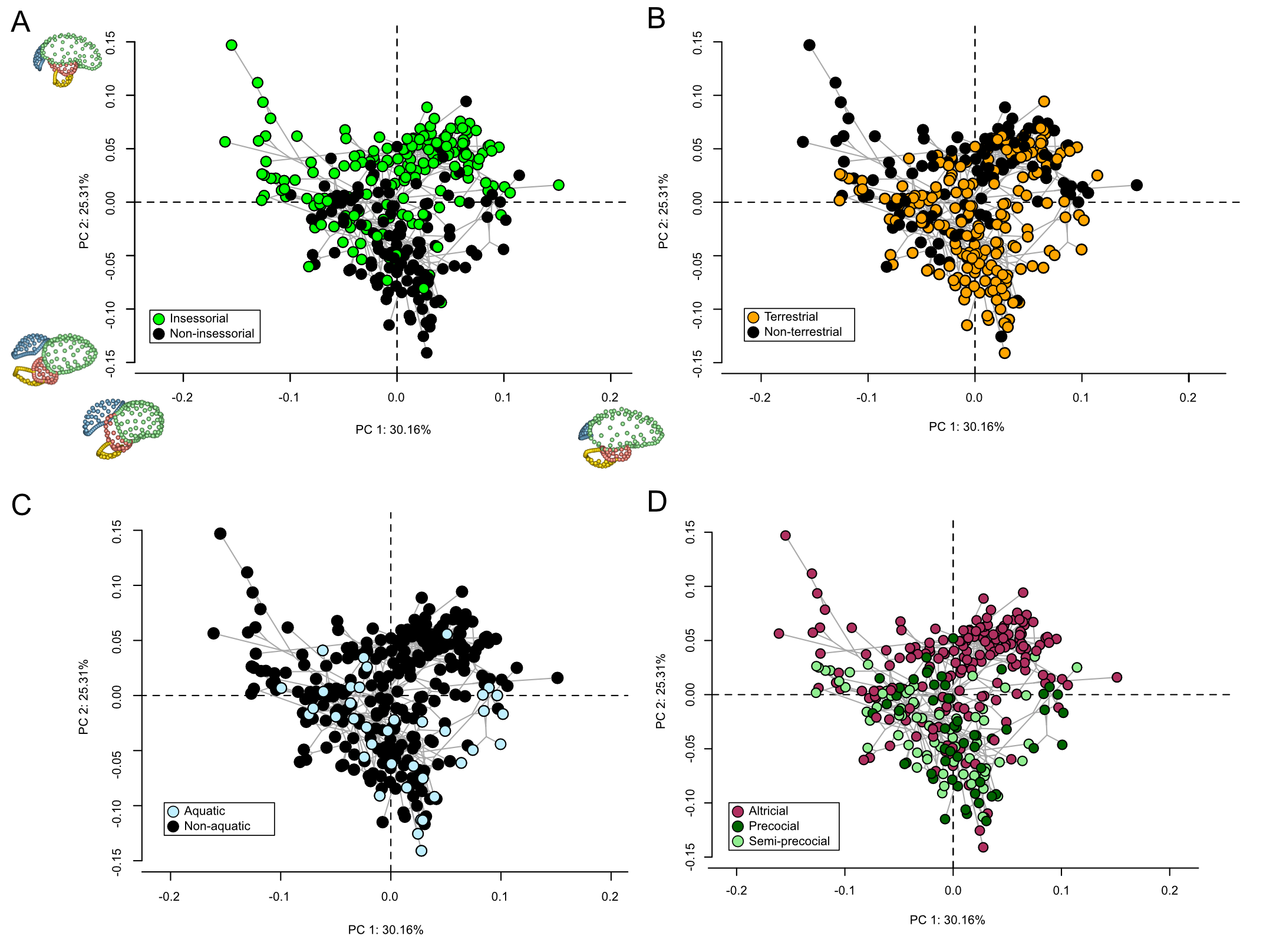

**Fig. S2:** Phylomorphospace of the first two principal component (PC) axes of extant taxa, with points colored by a) insessoriality, b) terrestrialty, c) aquatic lifestyle, and d) developmental mode.

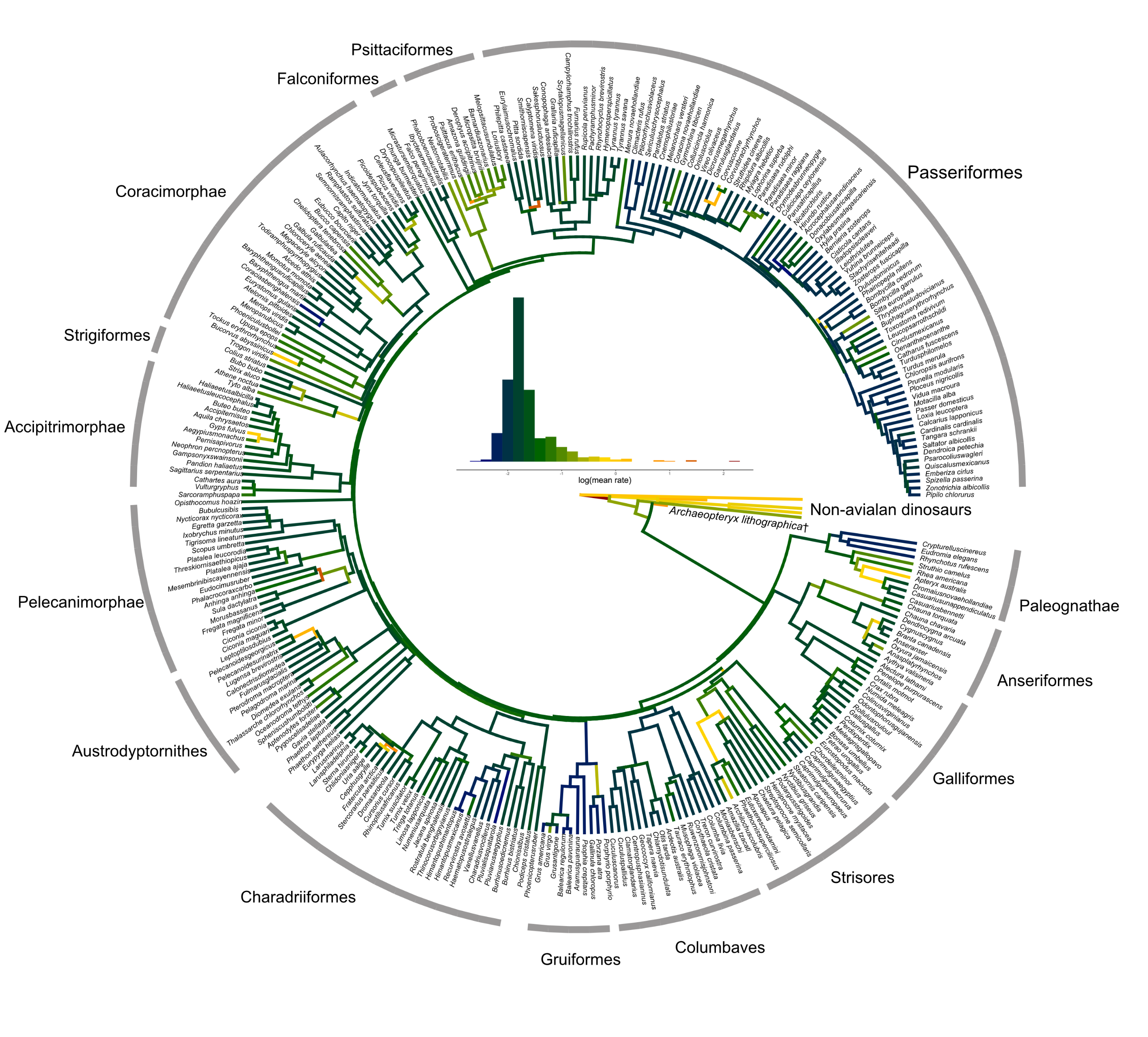

**Fig. S3:** Estimated rates of endocast shape evolution mapped onto time-calibrated phylogeny of birds under best-supported model of trait evolution (λ in all cases), in the whole endocast. Color gradient on branches indicates the rate of shape evolution as denoted by the inset histograms. Taxonomic naming from ref. ^3^.

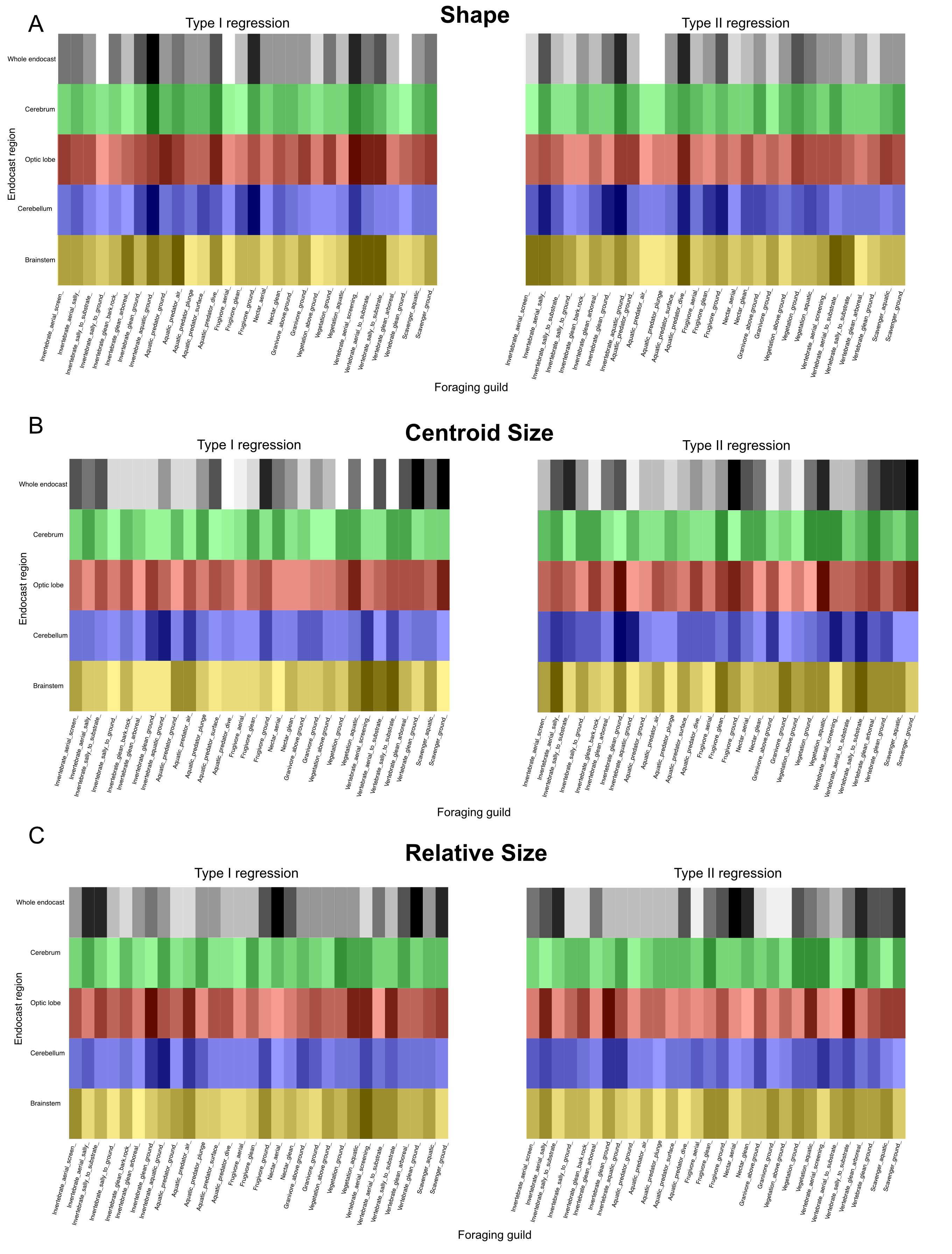

Fig. S4. Summary heatmap from ecomorphological analyses on foraging guild based on Z-scores, where darker gradient indicates greater values, thus relatively stronger association between endocast shape. Gradient is scaled to values along each row.

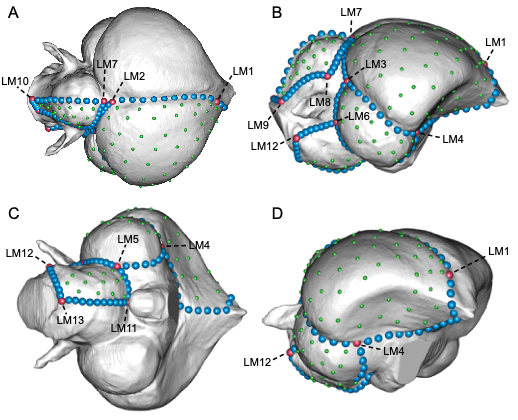

**Fig. S5:** Landmark scheme used in this study showing discrete (red), curve (blue), and surface (green) landmarks on the endocast of Gallus gallus (NMS Z1931.43) in A, dorsal; B, right lateral; C, ventral; and D, oblique views. The labels denote landmark numbers which correspond with Table S3.

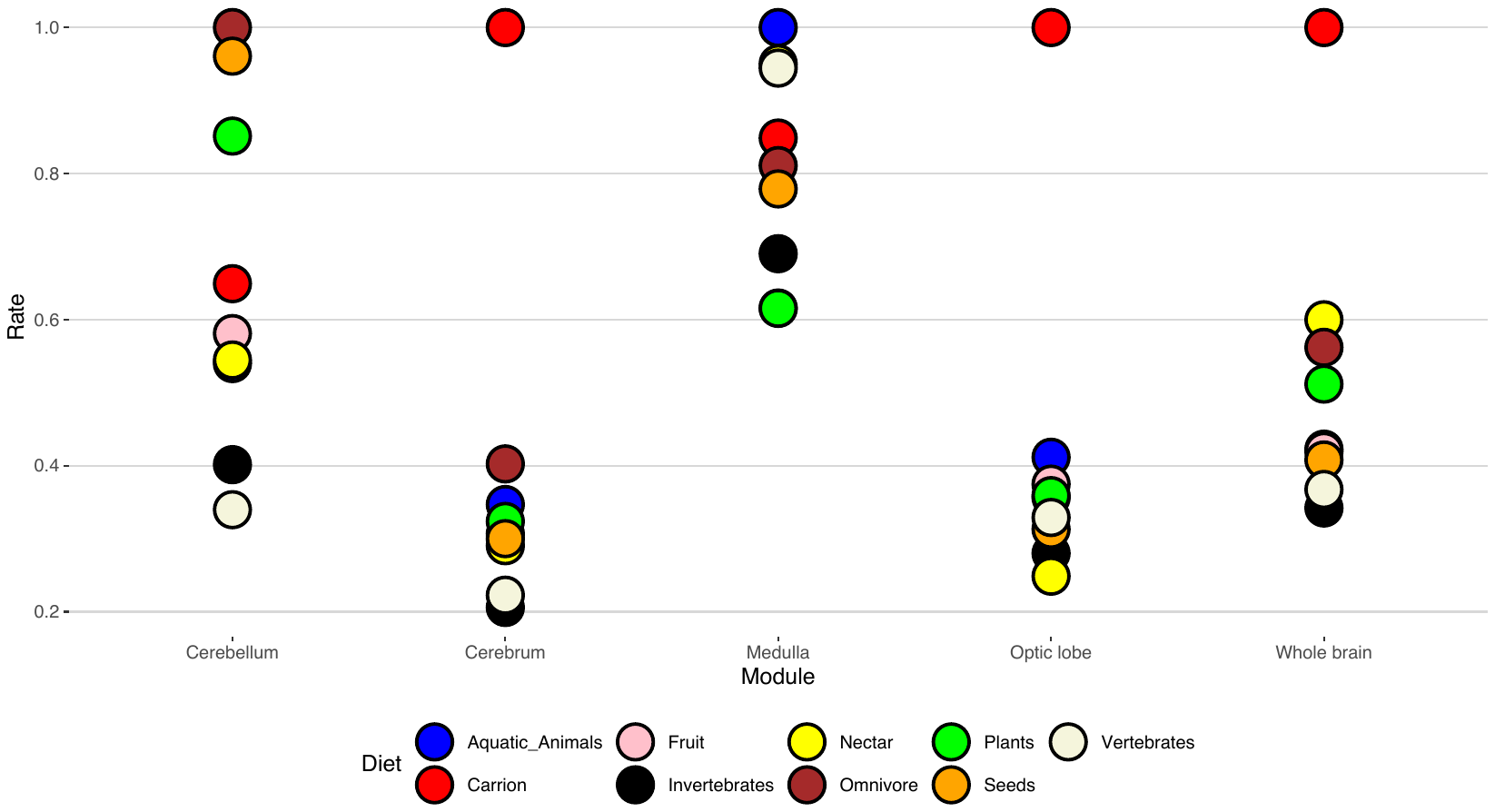

**Fig. S6:** Plot of standardized mean evolutionary rates for each endocast region colored by dietary category for the extant-only dataset.

**Table S1:** Results from phylogenetically corrected MANOVA on the extant-only dataset. For the Dietary Category column, a significance value is returned for the single categorical variable; the remaining columns indicate multivariate analyses, in names of variables found to be significantly correlated with the dependent morphological variable are provided. Statistical significance based on p-values of the covariation between ecological variables and the shape variable is indicated by symbols: 0 ‘***’ 0.001 ‘**’ 0.01 ‘*’ 0.05 ‘.’ 0.1 ‘ ’ 1

|  | | **Dietary category** | | | **Dietary guild** | | | | | | | **Locomotor ecology** | | | | | | |
| --- | --- | --- | --- | --- | --- | --- | --- | --- | --- | --- | --- | --- | --- | --- | --- | --- | --- | --- |
| **Dependent variable** | | **Rsq** | **Z** | ***p*** | **Variable** | **Type I** | | | **Type II** | | | **Variable** | **Type I** | | | **Type II** | | |
|  |  |  |  |  |  | **Rsq** | **Z** | ***p*** | **Rsq** | **Z** | ***p*** |  | **Rsq** | **Z** | ***p*** | **Rsq** | **Z** | ***p*** |
| **Whole brain**  K_mult_ = 0.62 | **Shape** | 0.04 | 2.14 | 0.02 * | **Frugivory** | 0.00434 | 0.69 | 0.26 | 0.00643 | 1.6480 | 0.055 . | **Volant** | 0.00111 | -1.3806 | 0.924 | 0.00124 | -1.1316 | 0.885 |
|  |  |  |  |  | **Nectarivory** | 0.0029 | 0.17 | 0.43 | 0.00407 | 0.9190 | 0.183 |  |  |  |  |  |  |  |
|  |  |  |  |  | **Granivory** | 0.004 | 0.94 | 0.17 | 0.00320 | 0.4416 | 0.331 | **Aquatic** | 0.00533 | 1.4335 | 0.067 . | 0.00554 | 1.4968 | 0.054 . |
|  |  |  |  |  | **Herbivory** | 0.0042 | 1.02 | 0.149 | 0.00428 | 1.1509 | 0.121 | **Soaring** | 0.00362 | 0.4627 | 0.337 | 0.00367 | 0.5060 | 0.317 |
|  |  |  |  |  | **Vertivory** | 0.0046 | 1.2 | 0.11 | 0.00402 | 0.9355 | 0.185 | **Arboreal** | 0.00466 | 1.1630 | 0.125 | 0.00368 | 0.6676 | 0.254 |
|  |  |  |  |  | **Invertivory** | 0.005 | 1.2 | 0.13 | 0.00379 | 0.8661 | 0.200 |  |  |  |  |  |  |  |
|  |  |  |  |  | **Aquatic predation** | 0.0045 | 1.16 | 0.131 | 0.01147 | 2.9355 | 0.004 ** | **Terrestrial** | 0.00779 | 2.3229 | 0.007 ** | 0.00734 | 2.1740 | 0.009 ** |
|  |  |  |  |  | **Scavenging** | 0.0054 | 1.44 | 0.07 . | 0.00663 | 1.9043 | 0.030 * |  |  |  |  |  |  |  |
|  | **Size** | 0.097 | 3.44 | 0.002 ** | **Frugivory** | 0.01288 | 1.6817 | 0.036 | 0.01905 | 2.2697 | 0.006 ** | **Volant** | 0.0009 | -0.249 | 0.601 | 0.00118 | -0.02744 | 0.514 |
|  |  |  |  |  | **Nectarivory** | 0.00003 | -1.503 | 0.92 | 0.00109 | -0.0390 | 0.535 |  |  |  |  |  |  |  |
|  |  |  |  |  | **Granivory** | 0.01823 | 1.7883 | 0.027 * | 0.00006 | -1.3256 | 0.889 | **Aquatic** | 0.00937 | 1.3164 | 0.099 . | 0.00795 | 1.20903 | 0.121 |
|  |  |  |  |  | **Herbivory** | 0.01096 | 1.4038 | 0.076 | 0.01170 | 1.6824 | 0.043 | **Soaring** | 0.02491 | 2.2518 | 0.006 ** | 0.02253 | 2.17849 | 0.008 ** |
|  |  |  |  |  | **Vertivory** | 0.02047 | 2.0554 | 0.01 ** | 0.07615 | 3.7765 | 0.001 *** | **Arboreal** | 0.02956 | 2.3611 | 0.004 ** | 0.02044 | 2.03865 | 0.011 * |
|  |  |  |  |  | **Invertivory** | 0.07827 | 3.8253 | 0.001 *** | 0.01681 | 2.0499 | 0.009 ** |  |  |  |  |  |  |  |
|  |  |  |  |  | **Aquatic predation** | 0.00135 | -0.032 | 0.509 | 0.00014 | -0.8252 | 0.783 | **Terrestrial** | 0.00942 | 1.2716 | 0.11 | 0.00390 | 0.66717 | 0.264 |
|  |  |  |  |  | **Scavenging** | 0.05931 | 3.4215 | 0.001 *** | 0.04782 | 3.2641 | 0.001 *** |  |  |  |  |  |  |  |
|  | **Relative size** | 0.087 | 2.82 | 0.003 ** | **Frugivory** | 0.00609 | 1.0008 | 0.164 | 0.00767 | 1.3115 | 0.091 . | **Volant** | 0.00059 | -0.301 | 0.629 | 0.00043 | -0.40348 | 0.660 |
|  |  |  |  |  | **Nectarivory** | 0.00001 | -1.777 | 0.956 | 0.0006 | -0.3168 | 0.626 |  |  |  |  |  |  |  |
|  |  |  |  |  | **Granivory** | 0.00357 | 0.5163 | 0.327 | 0.00081 | -0.3273 | 0.62 | **Aquatic** | 0.00919 | 1.2938 | 0.096 . | 0.00774 | 1.17185 | 0.125 |
|  |  |  |  |  | **Herbivory** | 0.0043 | 0.7333 | 0.261 | 0.02014 | 2.2232 | 0.009 ** | **Soaring** | 0.00645 | 1.0326 | 0.154 | 0.00525 | 0.88650 | 0.200 |
|  |  |  |  |  | **Vertivory** | 0.0242 | 2.1877 | 0.007 ** | 0.07792 | 3.6526 | 0.001 *** | **Arboreal** | 0.0194 | 1.9686 | 0.02 * | 0.01275 | 1.59645 | 0.054 . |
|  |  |  |  |  | **Invertivory** | 0.0562 | 3.2801 | 0.001 *** | 0.00215 | 0.3699 | 0.367 |  |  |  |  |  |  |  |
|  |  |  |  |  | **Aquatic predation** | 0.00039 | -0.653 | 0.732 | 0.00091 | -0.0989 | 0.55 | **Terrestrial** | 0.00997 | 1.3264 | 0.091 . | 0.00546 | 0.86242 | 0.214 |
|  |  |  |  |  | **Scavenging** | 0.02719 | 2.2436 | 0.013 * | 0.02383 | 2.3642 | 0.007 ** |  |  |  |  |  |  |  |
| **Cerebrum**  K_mult_ = 0.31 | **Shape** | 0.034 | 1.79 | 0.036 * | **Frugivory** | 0.0047 | 0.814 | 0.205 | 0.00483 | 0.8454 | 0.220 | **Volant** | 0.00191 | -0.1572 | 0.583 | 0.00215 | 0.0651 | 0.476 |
|  |  |  |  |  | **Nectarivory** | 0.0033 | 0.387 | 0.35 | 0.00286 | 0.2059 | 0.426 |  |  |  |  |  |  |  |
|  |  |  |  |  | **Granivory** | 0.00442 | 1.1376 | 0.129 | 0.00361 | 0.6900 | 0.244 | **Aquatic** | 0.00558 | 1.4493 | 0.069 . | 0.00571 | 1.4802 | 0.067 . |
|  |  |  |  |  | **Herbivory** | 0.00257 | -0.1283 | 0.555 | 0.00375 | 0.7237 | 0.244 | **Soaring** | 0.00335 | 0.2481 | 0.404 | 0.00332 | 0.2446 | 0.410 |
|  |  |  |  |  | **Vertivory** | 0.00582 | 1.6002 | 0.055 . | 0.00558 | 1.6250 | 0.051 . | **Arboreal** | 0.00386 | 0.6677 | 0.265 | 0.00374 | 0.6479 | 0.256 |
|  |  |  |  |  | **Invertivory** | 0.00553 | 1.2663 | 0.109 | 0.00342 | 0.5662 | 0.286 |  |  |  |  |  |  |  |
|  |  |  |  |  | **Aquatic predation** | 0.00530 | 1.4024 | 0.086 . | 0.00797 | 2.0770 | 0.019 * | **Terrestrial** | 0.00424 | 0.8129 | 0.236 | 0.00421 | 0.8149 | 0.226 |
|  |  |  |  |  | **Scavenging** | 0.00538 | 1.3552 | 0.089 . | 0.00654 | 1.7209 | 0.044 * |  |  |  |  |  |  |  |
|  | **Size** | 0.017 | -0.58 | 0.73 | **Frugivory** | 0.00087 | -0.2539 | 0.598 | 0.00220 | 0.25070 | 0.413 | **Volant** | 0.00028 | -0.66389 | 0.76 | 0.00030 | -0.52023 | 0.717 |
|  |  |  |  |  | **Nectarivory** | 0.00186 | 0.2216 | 0.43 | 0.00021 | -0.86133 | 0.796 |  |  |  |  |  |  |  |
|  |  |  |  |  | **Granivory** | 0.0006 | -0.3706 | 0.646 | 0.00155 | 0.02097 | 0.503 | **Aquatic** | 0.00005 | -1.3464 | 0.906 | 0.00000 | -2.43199 | 0.995 |
|  |  |  |  |  | **Herbivory** | 0.00508 | 0.7954 | 0.218 | 0.00167 | 0.05559 | 0.499 | **Soaring** | 0.00084 | -0.2929 | 0.638 | 0.00083 | -0.28659 | 0.642 |
|  |  |  |  |  | **Vertivory** | 0.00468 | 0.7828 | 0.227 | 0.00004 | -1.43412 | 0.912 | **Arboreal** | 0.00073 | -0.29209 | 0.63 | 0.00247 | 0.36807 | 0.367 |
|  |  |  |  |  | **Invertivory** | 0.00007 | -1.341 | 0.907 | 0.00196 | 0.15961 | 0.452 |  |  |  |  |  |  |  |
|  |  |  |  |  | **Aquatic predation** | 0.00195 | 0.18816 | 0.45 | 0.00249 | 0.35903 | 0.374 | **Terrestrial** | 0.00867 | 1.2032 | 0.117 | 0.01017 | 1.33165 | 0.087 . |
|  |  |  |  |  | **Scavenging** | 0.00476 | 0.8237 | 0.21 | 0.00024 | -0.83169 | 0.798 |  |  |  |  |  |  |  |
|  | **Relative size** | 0.016 | -0.63 | 0.75 | **Frugivory** | 0.00009 | -1.2466 | 0.881 | 0.00501 | 0.81931 | 0.236 | **Volant** | 0.00188 | 0.29465 | 0.413 | 0.00198 | 0.32114 | 0.372 |
|  |  |  |  |  | **Nectarivory** | 0.00009 | -1.1213 | 0.87 | 0.00213 | 0.27587 | 0.412 |  |  |  |  |  |  |  |
|  |  |  |  |  | **Granivory** | 0.00215 | 0.294 | 0.403 | 0.00242 | 0.32826 | 0.385 | **Aquatic** | 0 | -2.1359 | 0.983 | 0.00005 | -1.26195 | 0.888 |
|  |  |  |  |  | **Herbivory** | 0.0082 | 1.1636 | 0.118 | 0.00271 | 0.32632 | 0.382 | **Soaring** | 0.00217 | 0.22972 | 0.426 | 0.00215 | 0.22993 | 0.430 |
|  |  |  |  |  | **Vertivory** | 0.00084 | -0.2355 | 0.603 | 0.00081 | -0.30125 | 0.609 | **Arboreal** | 0.0001 | -1.138 | 0.863 | 0.00103 | -0.12729 | 0.563 |
|  |  |  |  |  | **Invertivory** | 0.00013 | -1.0687 | 0.835 | 0.00044 | -0.57092 | 0.718 |  |  |  |  |  |  |  |
|  |  |  |  |  | **Aquatic predation** | 0.00022 | -0.825 | 0.773 | 0.0004 | -0.60393 | 0.729 | **Terrestrial** | 0.00731 | 1.0513 | 0.158 | 0.00794 | 1.12274 | 0.143 |
|  |  |  |  |  | **Scavenging** | 0.00909 | 1.2735 | 0.1 | 0.00016 | -0.96918 | 0.822 |  |  |  |  |  |  |  |
| **Optic lobe**  K_mult_ = 0.53 | **Shape** | 0.045 | 2.68 | 0.002 ** | **Frugivory** | 0.00424 | 0.9269 | 0.178 | 0.00323 | 0.3128 | 0.368 | **Volant** | 0.0014 | -0.7020 | 0.759 | 0.00150 | -0.5866 | 0.720 |
|  |  |  |  |  | **Nectarivory** | 0.00379 | 0.6551 | 0.260 | 0.00380 | 0.6591 | 0.243 |  |  |  |  |  |  |  |
|  |  |  |  |  | **Granivory** | 0.01117 | 2.7828 | 0.006 ** | 0.00266 | -0.1229 | 0.533 | **Aquatic** | 0.0107 | 2.3696 | 0.013 * | 0.01061 | 2.3877 | 0.015 * |
|  |  |  |  |  | **Herbivory** | 0.00409 | 0.8773 | 0.201 | 0.00535 | 1.5272 | 0.061 . | **Soaring** | 0.0016 | -1.1051 | 0.858 | 0.00166 | -1.0384 | 0.844 |
|  |  |  |  |  | **Vertivory** | 0.00552 | 1.3493 | 0.090 . | 0.00310 | 0.2427 | 0.417 | **Arboreal** | 0.0057 | 1.4230 | 0.077 . | 0.00576 | 1.4694 | 0.069 . |
|  |  |  |  |  | **Invertivory** | 0.01048 | 2.9356 | 0.001 ** | 0.00647 | 1.7485 | 0.044 * |  |  |  |  |  |  |  |
|  |  |  |  |  | **Aquatic predation** | 0.00855 | 2.3149 | 0.011 * | 0.01042 | 2.4322 | 0.012 * | **Terrestrial** | 0.0076 | 2.2638 | 0.009 ** | 0.00822 | 2.4207 | 0.008 ** |
|  |  |  |  |  | **Scavenging** | 0.01299 | 2.5636 | 0.006 ** | 0.01186 | 2.8799 | 0.002 ** |  |  |  |  |  |  |  |
|  | **Size** | 0.1098 | 3.32 | 0.002 ** | **Frugivory** | 0.00918 | 1.2994 | 0.095 . | 0.01377 | 1.74410 | 0.039 * | **Volant** | 0.00092 | -0.2109 | 0.585 | 0.00149 | 0.09464 | 0.483 |
|  |  |  |  |  | **Nectarivory** | 0.00004 | -1.4027 | 0.905 | 0.00061 | -0.32457 | 0.628 |  |  |  |  |  |  |  |
|  |  |  |  |  | **Granivory** | 0.01577 | 1.6915 | 0.042 * | 0.01165 | 1.55909 | 0.053 . | **Aquatic** | 0.01315 | 1.5943 | 0.051 . | 0.01164 | 1.50163 | 0.059 . |
|  |  |  |  |  | **Herbivory** | 0.00377 | 0.6037 | 0.302 | 0.00719 | 1.16857 | 0.124 | **Soaring** | 0.01279 | 1.5582 | 0.058 . | 0.01204 | 1.50907 | 0.064 . |
|  |  |  |  |  | **Vertivory** | 0.00394 | 0.6638 | 0.271 | 0.01826 | 2.04699 | 0.011 * | **Arboreal** | 0.00689 | 1.13 | 0.134 | 0.00707 | 1.14052 | 0.123 |
|  |  |  |  |  | **Invertivory** | 0.05495 | 3.0235 | 0.001 *** | 0.03412 | 2.58082 | 0.003 ** |  |  |  |  |  |  |  |
|  |  |  |  |  | **Aquatic predation** | 0.02614 | 2.2865 | 0.004 ** | 0.01101 | 1.55384 | 0.054 . | **Terrestrial** | 0.00218 | 0.23179 | 0.429 | 0.00405 | 0.64122 | 0.272 |
|  |  |  |  |  | **Scavenging** | 0.07053 | 3.3549 | 0.001 *** | 0.01829 | 1.94520 | 0.021 * |  |  |  |  |  |  |  |
|  | **Relative size** | 0.119 | 3.9 | 0.001 *** | **Frugivory** | 0.00577 | 0.8836 | 0.2 | 0.00198 | 0.21121 | 0.432 | **Volant** | 0.00014 | -1.0703 | 0.857 | 0.00001 | -1.67732 | 0.947 |
|  |  |  |  |  | **Nectarivory** | 0.00171 | 0.0947 | 0.474 | 0.007 | 1.17376 | 0.119 |  |  |  |  |  |  |  |
|  |  |  |  |  | **Granivory** | 0.02452 | 2.1217 | 0.011 * | 0.01941 | 2.03438 | 0.014 * | **Aquatic** | 0.014 | 1.6326 | 0.043 * | 0.01178 | 1.48578 | 0.059 . |
|  |  |  |  |  | **Herbivory** | 0.00961 | 1.3299 | 0.096 . | 0.01155 | 1.58812 | 0.046 * | **Soaring** | 0.00752 | 1.131 | 0.143 | 0.00681 | 1.06510 | 0.162 |
|  |  |  |  |  | **Vertivory** | 0.01744 | 1.9104 | 0.024 * | 0.00578 | 1.03841 | 0.151 | **Arboreal** | 0.00731 | 1.1211 | 0.142 | 0.00809 | 1.17872 | 0.129 |
|  |  |  |  |  | **Invertivory** | 0.05326 | 3.066 | 0.001 *** | 0.05542 | 3.16172 | 0.001 *** |  |  |  |  |  |  |  |
|  |  |  |  |  | **Aquatic predation** | 0.05054 | 3.0149 | 0.001 *** | 0.0001 | -1.18251 | 0.867 | **Terrestrial** | 0.0028 | 0.38046 | 0.368 | 0.00489 | 0.77521 | 0.242 |
|  |  |  |  |  | **Scavenging** | 0.05993 | 3.1641 | 0.001 *** | 0.02276 | 2.234 | 0.007 ** |  |  |  |  |  |  |  |
| **Cerebellum**  K_mult_ = 0.14 | **Shape** | 0.041 | 1.59 | 0.058 . | **Frugivory** | 0.00480 | 0.95148 | 0.169 | 0.00761 | 1.8223 | 0.035 * | **Volant** | 0.00344 | 0.69949 | 0.239 | 0.00370 | 0.79733 | 0.207 |
|  |  |  |  |  | **Nectarivory** | 0.00343 | 0.5681 | 0.272 | 0.00625 | 1.3901 | 0.099 . |  |  |  |  |  |  |  |
|  |  |  |  |  | **Granivory** | 0.00497 | 1.0863 | 0.141 | 0.00309 | 0.2756 | 0.387 | **Aquatic** | 0.00601 | 1.2232 | 0.107 | 0.00631 | 1.31186 | 0.092 . |
|  |  |  |  |  | **Herbivory** | 0.00656 | 1.3916 | 0.083 . | 0.00152 | -1.0136 | 0.846 | **Soaring** | 0.00171 | -0.8031 | 0.783 | 0.00169 | -0.80802 | 0.789 |
|  |  |  |  |  | **Vertivory** | 0.00448 | 0.8335 | 0.205 | 0.00353 | 0.514 | 0.301 | **Arboreal** | 0.01094 | 2.0698 | 0.020 * | 0.00929 | 1.88923 | 0.029 * |
|  |  |  |  |  | **Invertivory** | 0.00274 | -0.0459 | 0.521 | 0.00797 | 1.9651 | 0.026 * |  |  |  |  |  |  |  |
|  |  |  |  |  | **Aquatic predation** | 0.00339 | 0.3726 | 0.361 | 0.03975 | 3.5213 | 0.001 *** | **Terrestrial** | 0.00476 | 0.8729 | 0.190 | 0.00407 | 0.60505 | 0.270 |
|  |  |  |  |  | **Scavenging** | 0.00212 | -0.1784 | 0.567 | 0.00199 | -0.4975 | 0.683 |  |  |  |  |  |  |  |
|  | **Size** | 0.026 | 0.15 | 0.443 | **Frugivory** | 0.00032 | -0.7135 | 0.751 | 0.00853 | 1.31986 | 0.094 . | **Volant** | 0.00227 | 0.40471 | 0.349 | 0.00210 | 0.27380 | 0.409 |
|  |  |  |  |  | **Nectarivory** | 0.00097 | -0.1702 | 0.574 | 0.00854 | 1.25339 | 0.107 |  |  |  |  |  |  |  |
|  |  |  |  |  | **Granivory** | 0.014 | 1.7342 | 0.037 * | 0.00800 | 1.25877 | 0.099 . | **Aquatic** | 0.00002 | -1.7007 | 0.949 | 0.00009 | -1.18895 | 0.868 |
|  |  |  |  |  | **Herbivory** | 0.00033 | -0.726 | 0.751 | 0.00051 | -0.49316 | 0.689 | **Soaring** | 0.0004 | -0.60826 | 0.716 | 0.00049 | -0.52308 | 0.696 |
|  |  |  |  |  | **Vertivory** | 0.00011 | -1.1606 | 0.861 | 0.00022 | -0.85813 | 0.773 | **Arboreal** | 0.00212 | 0.201 | 0.436 | 0.00046 | -0.58732 | 0.708 |
|  |  |  |  |  | **Invertivory** | 0.00141 | -0.02 | 0.519 | 0.01129 | 1.54312 | 0.059 . |  |  |  |  |  |  |  |
|  |  |  |  |  | **Aquatic predation** | 0.01029 | 1.4146 | 0.087 . | 0.01309 | 1.61693 | 0.051 . | **Terrestrial** | 0.01194 | 1.5144 | 0.065 . | 0.01041 | 1.37489 | 0.081 . |
|  |  |  |  |  | **Scavenging** | 0.00034 | -0.599 | 0.724 | 0.00190 | 0.17126 | 0.448 |  |  |  |  |  |  |  |
|  | **Relative size** | 0.028 | 0.25 | 0.424 | **Frugivory** | 0.00095 | -0.2107 | 0.597 | 0.00723 | 1.17491 | 0.129 | **Volant** | 0.00151 | 0.16957 | 0.446 | 0.00135 | 0.03829 | 0.497 |
|  |  |  |  |  | **Nectarivory** | 0.00015 | -0.9646 | 0.821 | 0.00657 | 1.03378 | 0.158 |  |  |  |  |  |  |  |
|  |  |  |  |  | **Granivory** | 0.0146 | 1.7884 | 0.03 * | 0.00851 | 1.3065 | 0.096 . | **Aquatic** | 0 | -2.622 | 0.999 | 0.00004 | -1.42559 | 0.911 |
|  |  |  |  |  | **Herbivory** | 0.00027 | -0.7908 | 0.779 | 0.00045 | -0.50463 | 0.692 | **Soaring** | 0.00116 | -0.0929 | 0.563 | 0.00132 | -0.02020 | 0.530 |
|  |  |  |  |  | **Vertivory** | 0.00067 | -0.4003 | 0.66 | 0.0001 | -1.13671 | 0.859 | **Arboreal** | 0.00137 | -0.0345 | 0.527 | 0.00010 | -1.15106 | 0.861 |
|  |  |  |  |  | **Invertivory** | 0.00269 | 0.3635 | 0.371 | 0.00977 | 1.40875 | 0.071 . |  |  |  |  |  |  |  |
|  |  |  |  |  | **Aquatic predation** | 0.00673 | 1.0823 | 0.145 | 0.00812 | 1.22296 | 0.099 . | **Terrestrial** | 0.01495 | 1.716 | 0.037 * | 0.01379 | 1.60744 | 0.048 * |
|  |  |  |  |  | **Scavenging** | 0.00117 | -0.0483 | 0.524 | 0.00219 | 0.26529 | 0.414 |  |  |  |  |  |  |  |
| **Brainstem**  K_mult_ = 0.27 | **Shape** | 0.034 | 1.22 | 0.112 | **Frugivory** | 0.00400 | 0.7282 | 0.232 | 0.00495 | 1.1385 | 0.129 | **Volant** | 0.00097 | -1.2857 | 0.898 | 0.00094 | -1.3764 | 0.913 |
|  |  |  |  |  | **Nectarivory** | 0.00594 | 1.3056 | 0.102 | 0.00346 | 0.4747 | 0.317 |  |  |  |  |  |  |  |
|  |  |  |  |  | **Granivory** | 0.00707 | 1.7154 | 0.040 * | 0.00688 | 1.6555 | 0.051 . | **Aquatic** | 0.00578 | 1.2366 | 0.107 | 0.00560 | 1.1914 | 0.118 |
|  |  |  |  |  | **Herbivory** | 0.00609 | 1.3756 | 0.093 . | 0.00251 | -0.1535 | 0.535 | **Soaring** | 0.00119 | -1.5466 | 0.934 | 0.00124 | -1.4631 | 0.921 |
|  |  |  |  |  | **Vertivory** | 0.01333 | 2.6539 | 0.005 ** | 0.00181 | -0.8528 | 0.801 | **Arboreal** | 0.00632 | 1.4611 | 0.069 . | 0.00527 | 1.1484 | 0.128 |
|  |  |  |  |  | **Invertivory** | 0.00603 | 1.5031 | 0.059 . | 0.00235 | -0.2280 | 0.589 |  |  |  |  |  |  |  |
|  |  |  |  |  | **Aquatic predation** | 0.00632 | 1.5091 | 0.066 . | 0.01086 | 2.2589 | 0.011 * | **Terrestrial** | 0.00357 | 0.5186 | 0.305 | 0.00277 | 0.027 | 0.508 |
|  |  |  |  |  | **Scavenging** | 0.00671 | 1.4829 | 0.068 . | 0.00351 | 0.4395 | 0.330 |  |  |  |  |  |  |  |
|  | **Size** | 0.058 | 1.72 | 0.047 * | **Frugivory** | 0.00001 | -1.7263 | 0.947 | 0.00079 | -0.35933 | 0.644 | **Volant** | 0.00218 | 0.33809 | 0.397 | 0.00197 | 0.26982 | 0.419 |
|  |  |  |  |  | **Nectarivory** | 0.00313 | 0.5677 | 0.308 | 0.00407 | 0.68477 | 0.265 |  |  |  |  |  |  |  |
|  |  |  |  |  | **Granivory** | 0.00135 | -0.0387 | 0.519 | 0.0056 | 0.92917 | 0.178 | **Aquatic** | 0.01178 | 1.529 | 0.059 . | 0.01011 | 1.40798 | 0.076 . |
|  |  |  |  |  | **Herbivory** | 0.05028 | 2.9632 | 0.002 ** | 0.02653 | 2.28462 | 0.004 ** | **Soaring** | 0.0046 | 0.76521 | 0.232 | 0.00569 | 0.92155 | 0.186 |
|  |  |  |  |  | **Vertivory** | 0.04174 | 2.9372 | 0.002 ** | 0.00211 | 0.24721 | 0.424 | **Arboreal** | 0.01627 | 1.813 | 0.027 * | 0.01361 | 1.67009 | 0.038 * |
|  |  |  |  |  | **Invertivory** | 0.01197 | 1.5795 | 0.056 . | 0.00817 | 1.21607 | 0.11 |  |  |  |  |  |  |  |
|  |  |  |  |  | **Aquatic predation** | 0.01488 | 1.6696 | 0.041 * | 0.00046 | -0.53816 | 0.689 | **Terrestrial** | 0.00233 | 0.32225 | 0.403 | 0.00069 | -0.38232 | 0.662 |
|  |  |  |  |  | **Scavenging** | 0.00041 | -0.5678 | 0.708 | 0.004 | 0.62759 | 0.286 |  |  |  |  |  |  |  |
|  | **Relative size** | 0.066 | 2.12 | 0.021 * | **Frugivory** | 0.00057 | -0.4676 | 0.68 | 0.00398 | 0.627 | 0.274 | **Volant** | 0.00065 | -0.32097 | 0.625 | 0.00052 | -0.48759 | 0.690 |
|  |  |  |  |  | **Nectarivory** | 0.00081 | -0.2387 | 0.598 | 0.0014 | 0.00842 | 0.521 |  |  |  |  |  |  |  |
|  |  |  |  |  | **Granivory** | 0.00062 | -0.4436 | 0.664 | 0.00461 | 0.813 | 0.217 | **Aquatic** | 0.01534 | 1.7358 | 0.037 * | 0.01321 | 1.61523 | 0.044 * |
|  |  |  |  |  | **Herbivory** | 0.05553 | 3.0618 | 0.001 *** | 0.02849 | 2.399 | 0.003 ** | **Soaring** | 0.00319 | 0.48662 | 0.317 | 0.00421 | 0.68735 | 0.253 |
|  |  |  |  |  | **Vertivory** | 0.03027 | 2.4528 | 0.002 ** | 0.0001 | -1.073 | 0.842 | **Arboreal** | 0.02556 | 2.2682 | 0.01 ** | 0.02223 | 2.12135 | 0.013 * |
|  |  |  |  |  | **Invertivory** | 0.01783 | 1.9806 | 0.015 * | 0.00493 | 0.795 | 0.225 |  |  |  |  |  |  |  |
|  |  |  |  |  | **Aquatic predation** | 0.0084 | 1.165 | 0.123 | 0.0074 | 1.107 | 0.135 | **Terrestrial** | 0.00184 | 0.17533 | 0.448 | 0.00019 | -0.89983 | 0.806 |
|  |  |  |  |  | **Scavenging** | 0.00167 | 0.1327 | 0.457 | 0.00458 | 0.731 | 0.25 |  |  |  |  |  |  |  |

**Table S2:** Results from phylogenetically corrected MANOVA on the extant-only dataset for ecomorphological analyses of foraging guild. Statistical significance based on p-values of the covariation between ecological variables and the shape variable is indicated by symbols: 0 ‘***’ 0.001 ‘**’ 0.01 ‘*’ 0.05 ‘.’ 0.1 ‘ ’ 1

|  | | **Foraging guild** | | | | | | |
| --- | --- | --- | --- | --- | --- | --- | --- | --- |
| **Dependent variable** | | **Variable** | **Type I** | | | **Type II** | | |
|  |  |  | **Rsq** | **Z** | ***p*** | **Rsq** | **Z** | ***p*** |
| **Whole brain**  K_mult_ = 0.62 | **Shape** | **Invertebrate aerial screen** | 0.00492 | 1.2068 | 0.112 | 0.00211 | -0.515 | 0.706 |
|  |  | **Invertebrate aerial sally** | 0.00511 | 1.4972 | 0.072 . | 0.0055 | 1.7431 | 0.041 * |
|  |  | **Invertebrate sally to substrate** | 0.00357 | 0.7395 | 0.240 | 0.00267 | -0.0263 | 0.502 |
|  |  | **Invertebrate sally to ground** | 0.00144 | -1.2456 | 0.896 | 0.00194 | -0.693 | 0.75 |
|  |  | **Invertebrate glean bark.rock** | 0.00507 | 1.2927 | 0.100 . | 0.00379 | 0.7757 | 0.238 |
|  |  | **Invertebrate glean arboreal** | 0.00282 | 0.0893 | 0.453 | 0.00267 | 0.0324 | 0.477 |
|  |  | **Invertebrate glean ground** | 0.00676 | 1.9221 | 0.026 * | 0.0055 | 1.6219 | 0.051 . |
|  |  | **Invertebrate aquatic ground** | 0.01618 | 2.9366 | 0.004 ** | 0.0075 | 2.3708 | 0.009 ** |
|  |  | **Aquatic predator ground** | 0.00384 | 0.8428 | 0.182 | 0.00318 | 0.3763 | 0.37 |
|  |  | **Aquatic predator air** | 0.00605 | 1.6306 | 0.056 . | 0.00108 | -2.1976 | 0.99 |
|  |  | **Aquatic predator plunge** | 0.00425 | 0.8858 | 0.189 | 0.0011 | -2.0016 | 0.979 |
|  |  | **Aquatic predator surface** | 0.00575 | 1.1246 | 0.133 | 0.00411 | 0.7135 | 0.254 |
|  |  | **Aquatic predator dive** | 0.00785 | 2.2898 | 0.013 * | 0.00793 | 2.3417 | 0.011 * |
|  |  | **Frugivore aerial** | 0.00154 | -1.0784 | 0.871 | 0.00245 | -0.1179 | 0.543 |
|  |  | **Frugivore glean** | 0.00313 | 0.2972 | 0.374 | 0.00385 | 0.8333 | 0.217 |
|  |  | **Frugivore ground** | 0.01175 | 2.8300 | 0.005 ** | 0.00623 | 1.753 | 0.036 * |
|  |  | **Nectar aerial** | 0.00395 | 0.8223 | 0.210 | 0.00199 | -0.3267 | 0.633 |
|  |  | **Nectar glean** | 0.00369 | 0.6953 | 0.241 | 0.00286 | 0.2667 | 0.393 |
|  |  | **Granivore above.ground** | 0.00440 | 1.0900 | 0.139 | 0.00365 | 0.8951 | 0.191 |
|  |  | **Granivore ground** | 0.00436 | 0.7879 | 0.217 | 0.00199 | -0.778 | 0.775 |
|  |  | **Vegetation above.ground** | 0.00286 | 0.0679 | 0.482 | 0.00368 | 0.7964 | 0.216 |
|  |  | **Vegetation ground** | 0.00550 | 1.6061 | 0.060 . | 0.00713 | 2.2364 | 0.013 * |
|  |  | **Vegetation aquatic** | 0.00344 | 0.6368 | 0.259 | 0.00408 | 1.1156 | 0.14 |
|  |  | **Vertebrate aerial screening** | 0.01330 | 2.3398 | 0.009 ** | 0.0038 | 0.8536 | 0.204 |
|  |  | **Vertebrate aerial to substrate** | 0.00527 | 1.3676 | 0.084 . | 0.00406 | 0.9789 | 0.172 |
|  |  | **Vertebrate sally to substrate** | 0.00611 | 1.8223 | 0.034 * | 0.00289 | 0.3126 | 0.38 |
|  |  | **Vertebrate glean arboreal** | 0.00373 | 0.4985 | 0.306 | 0.00366 | 0.6345 | 0.263 |
|  |  | **Vertebrate glean ground** | 0.00174 | -1.1950 | 0.885 | 0.00214 | -0.5922 | 0.714 |
|  |  | **Scavenger aquatic** | 0.00508 | 1.0749 | 0.134 | 0.00403 | 0.8442 | 0.203 |
|  |  | **Scavenger ground** | 0.00516 | 1.4314 | 0.077 . | 0.00396 | 0.9785 | 0.174 |
|  | **Size** | **Invertebrate aerial screen** | 0.03192 | 2.5073 | 0.006 ** | 0.00105 | -0.11036 | 0.553 |
|  |  | **Invertebrate aerial sally** | 0.01974 | 1.8791 | 0.022 * | 0.00801 | 1.23366 | 0.103 |
|  |  | **Invertebrate sally to substrate** | 0.02797 | 2.3395 | 0.003 ** | 0.01273 | 1.76609 | 0.026 * |
|  |  | **Invertebrate sally to ground** | 0.0005 | -0.51285 | 0.694 | 0.00191 | 0.27541 | 0.417 |
|  |  | **Invertebrate glean bark.rock** | 0.00074 | -0.35695 | 0.641 | 0.00004 | -1.33844 | 0.896 |
|  |  | **Invertebrate glean arboreal** | 0.00104 | -0.21635 | 0.592 | 0.00393 | 0.7458 | 0.247 |
|  |  | **Invertebrate glean ground** | 0.00123 | -0.065981 | 0.529 | 0.0093 | 1.40345 | 0.084 . |
|  |  | **Invertebrate aquatic ground** | 0.0042 | 0.69953 | 0.268 | 0.00002 | -1.62278 | 0.937 |
|  |  | **Aquatic predator ground** | 0.00042 | -0.62099 | 0.733 | 0.00063 | -0.34942 | 0.627 |
|  |  | **Aquatic predator air** | 0.00076 | -0.30573 | 0.63 | 0.00056 | -0.38381 | 0.648 |
|  |  | **Aquatic predator plunge** | 0.00616 | 0.97216 | 0.172 | 0.00027 | -0.68996 | 0.753 |
|  |  | **Aquatic predator surface** | 0.01967 | 1.9854 | 0.016 * | 0.00058 | -0.36457 | 0.636 |
|  |  | **Aquatic predator dive** | 0.00002 | -1.6101 | 0.936 | 0.00224 | 0.42082 | 0.36 |
|  |  | **Frugivore aerial** | 0.00012 | -1.1461 | 0.865 | 0.00041 | -0.44178 | 0.668 |
|  |  | **Frugivore glean** | 0.00057 | -0.47955 | 0.683 | 0.00268 | 0.47421 | 0.326 |
|  |  | **Frugivore ground** | 0.04871 | 3.0777 | 0.001 *** | 0.02515 | 2.43874 | 0.004 ** |
|  |  | **Nectar aerial** | 0.00933 | 1.3058 | 0.1 | 0.01037 | 1.50017 | 0.072 . |
|  |  | **Nectar glean** | 0.00537 | 0.87244 | 0.209 | 0.00538 | 1.04636 | 0.158 |
|  |  | **Granivore above.ground** | 0.0012 | -0.072376 | 0.535 | 0.00003 | -1.51361 | 0.926 |
|  |  | **Granivore ground** | 0.00532 | 0.8498 | 0.216 | 0.00058 | -0.40141 | 0.662 |
|  |  | **Vegetation above.ground** | 0.002 | 0.14349 | 0.462 | 0.00003 | -1.44924 | 0.916 |
|  |  | **Vegetation ground** | 0.00002 | -1.6792 | 0.944 | 0.00451 | 0.89326 | 0.201 |
|  |  | **Vegetation aquatic** | 0.00963 | 1.3302 | 0.103 | 0.01575 | 1.92691 | 0.021 * |
|  |  | **Vertebrate aerial screening** | 0.00000036 | -1.959 | 0.975 | 0.00077 | -0.22782 | 0.591 |
|  |  | **Vertebrate aerial to substrate** | 0.02008 | 1.9549 | 0.017 * | 0.00038 | -0.547 | 0.691 |
|  |  | **Vertebrate sally to substrate** | 0.00001 | -1.6822 | 0.944 | 0.00173 | 0.22292 | 0.431 |
|  |  | **Vertebrate glean arboreal** | 0.02144 | 2.0099 | 0.013 * | 0.00977 | 1.56324 | 0.052 |
|  |  | **Vertebrate glean ground** | 0.07535 | 3.6971 | 0.001 *** | 0.01807 | 2.09545 | 0.013 * |
|  |  | **Scavenger aquatic** | 0.01856 | 1.9185 | 0.019 * | 0.01435 | 1.8926 | 0.021 * |
|  |  | **Scavenger ground** | 0.07834 | 3.6176 | 0.001 *** | 0.02196 | 2.32679 | 0.004 ** |
|  | **Relative size** | **Invertebrate aerial screen** | 0.00292 | 0.43681 | 0.354 | 0.00705 | 1.2086 | 0.118 |
|  |  | **Invertebrate aerial sally** | 0.05415 | 2.9156 | 0.002 ** | 0.01834 | 2.0703 | 0.009 ** |
|  |  | **Invertebrate sally to substrate** | 0.04154 | 2.6942 | 0.003 ** | 0.03474 | 2.7613 | 0.003 ** |
|  |  | **Invertebrate sally to ground** | 0.00117 | -0.013475 | 0.518 | 0.00014 | -0.9702 | 0.817 |
|  |  | **Invertebrate glean bark.rock** | 0.00025 | -0.77 | 0.77 | 0.00022 | -0.7481 | 0.76 |
|  |  | **Invertebrate glean arboreal** | 0.00257 | 0.38018 | 0.379 | 0.00508 | 1.0172 | 0.16 |
|  |  | **Invertebrate glean ground** | 0.00779 | 1.1562 | 0.138 | 0.00165 | 0.165 | 0.466 |
|  |  | **Invertebrate aquatic ground** | 0.00289 | 0.47142 | 0.336 | 0.00116 | -0.0164 | 0.518 |
|  |  | **Aquatic predator ground** | 0.00027 | -0.72723 | 0.767 | 0.00103 | -0.0489 | 0.534 |
|  |  | **Aquatic predator air** | 0.00048 | -0.4097 | 0.646 | 0.0005 | -0.4218 | 0.663 |
|  |  | **Aquatic predator plunge** | 0.00512 | 0.86097 | 0.192 | 0.00088 | -0.0199 | 0.523 |
|  |  | **Aquatic predator surface** | 0.00393 | 0.62847 | 0.276 | 0.00082 | -0.116 | 0.562 |
|  |  | **Aquatic predator dive** | 0.00212 | 0.27624 | 0.409 | 0.00889 | 1.5456 | 0.054 . |
|  |  | **Frugivore aerial** | 0.00173 | 0.15889 | 0.441 | 0.00001 | -1.517 | 0.931 |
|  |  | **Frugivore glean** | 0.00192 | 0.13857 | 0.462 | 0.00789 | 1.3628 | 0.089 . |
|  |  | **Frugivore ground** | 0.01752 | 1.9488 | 0.018 * | 0.01081 | 1.68 | 0.039 * |
|  |  | **Nectar aerial** | 0.12642 | 3.7091 | 0.001 *** | 0.07515 | 3.5048 | 0.001 *** |
|  |  | **Nectar glean** | 0.02344 | 2.1318 | 0.013 * | 0.03111 | 2.6505 | 0.002 ** |
|  |  | **Granivore above.ground** | 0.00262 | 0.37581 | 0.367 | 0.00035 | -0.5816 | 0.716 |
|  |  | **Granivore ground** | 0.00647 | 1.0092 | 0.168 | 0.00009 | -1.1981 | 0.867 |
|  |  | **Vegetation above.ground** | 0.00439 | 0.76855 | 0.243 | 0.00004 | -1.3182 | 0.891 |
|  |  | **Vegetation ground** | 0.00322 | 0.52765 | 0.306 | 0.00889 | 1.5266 | 0.062 . |
|  |  | **Vegetation aquatic** | 0.00344 | 0.58097 | 0.297 | 0.00502 | 1.0441 | 0.145 |
|  |  | **Vertebrate aerial screening** | 0.00009 | -0.93528 | 0.817 | 0.00202 | 0.3981 | 0.363 |
|  |  | **Vertebrate aerial to substrate** | 0.00694 | 1.0963 | 0.142 | 0.00054 | -0.317 | 0.636 |
|  |  | **Vertebrate sally to substrate** | 0.00156 | 0.047266 | 0.5 | 0.00794 | 1.3495 | 0.091 . |
|  |  | **Vertebrate glean arboreal** | 0.02676 | 2.2015 | 0.01 ** | 0.01775 | 2.2241 | 0.005 ** |
|  |  | **Vertebrate glean ground** | 0.05612 | 3.1739 | 0.001 *** | 0.01272 | 1.8822 | 0.02 * |
|  |  | **Scavenger aquatic** | 0.00466 | 0.80743 | 0.214 | 0.00885 | 1.5235 | 0.056 . |
|  |  | **Scavenger ground** | 0.05217 | 2.952 | 0.001 *** | 0.02116 | 2.3845 | 0.003 ** |
| **Cerebrum**  K_mult_ = 0.31 | **Shape** | **Invertebrate aerial screen** | 0.00311 | 0.3427 | 0.366 | 0.00134 | -1.569 | 0.941 |
|  |  | **Invertebrate aerial sally** | 0.00359 | 0.6482 | 0.261 | 0.00465 | 1.1388 | 0.127 |
|  |  | **Invertebrate sally to substrate** | 0.00284 | 0.2616 | 0.402 | 0.00211 | -0.6032 | 0.729 |
|  |  | **Invertebrate sally to ground** | 0.00217 | -0.2992 | 0.617 | 0.00199 | -0.5765 | 0.709 |
|  |  | **Invertebrate glean bark.rock** | 0.00290 | 0.1125 | 0.454 | 0.00387 | 0.7089 | 0.25 |
|  |  | **Invertebrate glean arboreal** | 0.00204 | -0.6764 | 0.750 | 0.00263 | -0.0379 | 0.509 |
|  |  | **Invertebrate glean ground** | 0.00392 | 0.6155 | 0.280 | 0.00328 | 0.3559 | 0.354 |
|  |  | **Invertebrate aquatic ground** | 0.01203 | 2.5170 | 0.006 ** | 0.00636 | 1.8385 | 0.028 * |
|  |  | **Aquatic predator ground** | 0.00439 | 0.9750 | 0.158 | 0.00413 | 0.815 | 0.207 |
|  |  | **Aquatic predator air** | 0.00578 | 1.4988 | 0.076 . | 0.00162 | -1.0239 | 0.844 |
|  |  | **Aquatic predator plunge** | 0.00480 | 1.034 | 0.152 | 0.00147 | -1.1621 | 0.874 |
|  |  | **Aquatic predator surface** | 0.00614 | 1.1518 | 0.125 | 0.0042 | 0.6344 | 0.272 |
|  |  | **Aquatic predator dive** | 0.00839 | 2.2532 | 0.011 * | 0.005 | 1.355 | 0.093 . |
|  |  | **Frugivore aerial** | 0.00186 | -0.5978 | 0.720 | 0.0032 | 0.3893 | 0.361 |
|  |  | **Frugivore glean** | 0.00174 | -1.1967 | 0.890 | 0.00266 | -0.1474 | 0.552 |
|  |  | **Frugivore ground** | 0.00754 | 1.6528 | 0.043 * | 0.00373 | 0.485 | 0.318 |
|  |  | **Nectar aerial** | 0.00343 | 0.5417 | 0.281 | 0.0023 | -0.0075 | 0.508 |
|  |  | **Nectar glean** | 0.00383 | 0.7744 | 0.229 | 0.00273 | 0.1693 | 0.43 |
|  |  | **Granivore above.ground** | 0.00441 | 1.0188 | 0.170 | 0.00476 | 1.3829 | 0.085 . |
|  |  | **Granivore ground** | 0.00379 | 0.3920 | 0.352 | 0.00182 | -1.0086 | 0.834 |
|  |  | **Vegetation above.ground** | 0.00282 | -0.0081 | 0.508 | 0.00333 | 0.446 | 0.331 |
|  |  | **Vegetation ground** | 0.00327 | 0.3987 | 0.353 | 0.00595 | 1.5215 | 0.06 . |
|  |  | **Vegetation aquatic** | 0.00330 | 0.5344 | 0.307 | 0.00381 | 0.8602 | 0.194 |
|  |  | **Vertebrate aerial screening** | 0.01235 | 2.1410 | 0.018 * | 0.00279 | 0.2761 | 0.383 |
|  |  | **Vertebrate aerial to substrate** | 0.00616 | 1.5148 | 0.066 . | 0.00453 | 1.0836 | 0.146 |
|  |  | **Vertebrate sally to substrate** | 0.00445 | 1.0887 | 0.142 | 0.00288 | 0.2328 | 0.413 |
|  |  | **Vertebrate glean arboreal** | 0.00268 | -0.1808 | 0.573 | 0.00259 | -0.1709 | 0.577 |
|  |  | **Vertebrate glean ground** | 0.00152 | -1.4537 | 0.928 | 0.00284 | 0.1426 | 0.44 |
|  |  | **Scavenger aquatic** | 0.00568 | 1.1984 | 0.117 | 0.00447 | 0.9842 | 0.16 |
|  |  | **Scavenger ground** | 0.00532 | 1.4298 | 0.069 . | 0.00456 | 1.1806 | 0.122 |
|  | **Size** | **Invertebrate aerial screen** | 0.00182 | 0.23451 | 0.416 | 0.0025 | 0.38935 | 0.359 |
|  |  | **Invertebrate aerial sally** | 0.01618 | 1.7614 | 0.039 * | 0.00678 | 1.1438 | 0.143 |
|  |  | **Invertebrate sally to substrate** | 0.00232 | 0.39006 | 0.351 | 0.00009 | -1.14417 | 0.87 |
|  |  | **Invertebrate sally to ground** | 0.00002 | -1.5731 | 0.933 | 0.00399 | 0.68111 | 0.254 |
|  |  | **Invertebrate glean bark.rock** | 0.00636 | 0.98755 | 0.163 | 0.00741 | 1.13522 | 0.121 |
|  |  | **Invertebrate glean arboreal** | 0.00188 | 0.19851 | 0.441 | 0.0003 | -0.77694 | 0.763 |
|  |  | **Invertebrate glean ground** | 0.00044 | -0.66777 | 0.743 | 0.00001 | -1.82457 | 0.96 |
|  |  | **Invertebrate aquatic ground** | 0.00031 | -0.67714 | 0.743 | 0.00173 | 0.13877 | 0.449 |
|  |  | **Aquatic predator ground** | 0.00618 | 1.0652 | 0.133 | 0.00016 | -0.94592 | 0.815 |
|  |  | **Aquatic predator air** | 0.0002 | -0.85933 | 0.803 | 0.00021 | -0.84173 | 0.793 |
|  |  | **Aquatic predator plunge** | 0.00891 | 1.2747 | 0.095 . | 0.00266 | 0.48206 | 0.33 |
|  |  | **Aquatic predator surface** | 0.00002 | -1.5342 | 0.93 | 0.00108 | -0.06874 | 0.523 |
|  |  | **Aquatic predator dive** | 0.00018 | -0.95352 | 0.827 | 0.00036 | -0.62889 | 0.733 |
|  |  | **Frugivore aerial** | 0.00043 | -0.47967 | 0.689 | 0.00011 | -1.06015 | 0.844 |
|  |  | **Frugivore glean** | 0.01061 | 1.3498 | 0.084 . | 0.01143 | 1.50471 | 0.065 . |
|  |  | **Frugivore ground** | 0.00116 | -0.11538 | 0.566 | 0 | -1.88855 | 0.972 |
|  |  | **Nectar aerial** | 0.00976 | 1.3626 | 0.07 . | 0.00415 | 0.77934 | 0.214 |
|  |  | **Nectar glean** | 0.00102 | -0.093318 | 0.551 | 0.00483 | 0.85113 | 0.199 |
|  |  | **Granivore above.ground** | 0.00359 | 0.61256 | 0.28 | 0.00035 | -0.62384 | 0.724 |
|  |  | **Granivore ground** | 0.00001 | -1.8873 | 0.964 | 0.00111 | -0.1458 | 0.572 |
|  |  | **Vegetation above.ground** | 0.00005 | -1.3341 | 0.9 | 0.00001 | -1.64485 | 0.94 |
|  |  | **Vegetation ground** | 0.0132 | 1.5193 | 0.058 . | 0.01586 | 1.70606 | 0.043 * |
|  |  | **Vegetation aquatic** | 0.01241 | 1.5437 | 0.063 . | 0.01292 | 1.63825 | 0.046 * |
|  |  | **Vertebrate aerial screening** | 0.00036 | -0.46465 | 0.685 | 0.00897 | 1.2611 | 0.098 . |
|  |  | **Vertebrate aerial to substrate** | 0.00054 | -0.34341 | 0.637 | 0.00067 | -0.31597 | 0.626 |
|  |  | **Vertebrate sally to substrate** | 0.01607 | 1.8597 | 0.021 * | 0.00244 | 0.33227 | 0.394 |
|  |  | **Vertebrate glean arboreal** | 0.00994 | 1.2956 | 0.1 | 0.00898 | 1.29941 | 0.096 . |
|  |  | **Vertebrate glean ground** | 0.00082 | -0.29956 | 0.62 | 0.00322 | 0.48472 | 0.329 |
|  |  | **Scavenger aquatic** | 0.00286 | 0.51175 | 0.32 | 0.00054 | -0.43326 | 0.664 |
|  |  | **Scavenger ground** | 0.00343 | 0.57444 | 0.311 | 0.0049 | 0.82822 | 0.22 |
|  | **Relative size** | **Invertebrate aerial screen** | 0.00091 | -0.12648 | 0.565 | 0.00074 | -0.26408 | 0.615 |
|  |  | **Invertebrate aerial sally** | 0.01117 | 1.4591 | 0.065 . | 0.00004 | -1.41139 | 0.91 |
|  |  | **Invertebrate sally to substrate** | 0.00152 | 0.1219 | 0.481 | 0.0007 | -0.37724 | 0.646 |
|  |  | **Invertebrate sally to ground** | 0.00005 | -1.3656 | 0.911 | 0.00369 | 0.58809 | 0.287 |
|  |  | **Invertebrate glean bark.rock** | 0.0054 | 0.85146 | 0.202 | 0.00477 | 0.80205 | 0.228 |
|  |  | **Invertebrate glean arboreal** | 0.00058 | -0.46464 | 0.672 | 0.00036 | -0.64212 | 0.729 |
|  |  | **Invertebrate glean ground** | 0.00006 | -1.3671 | 0.908 | 0.0016 | 0.05864 | 0.496 |
|  |  | **Invertebrate aquatic ground** | 0.01183 | 1.4656 | 0.06 . | 0.00653 | 1.02329 | 0.157 |
|  |  | **Aquatic predator ground** | 0.0004 | -0.50347 | 0.692 | 0.00004 | -1.40988 | 0.91 |
|  |  | **Aquatic predator air** | 0.00129 | -0.0046293 | 0.514 | 0.00145 | 0.09309 | 0.478 |
|  |  | **Aquatic predator plunge** | 0.01173 | 1.4835 | 0.066 . | 0.00199 | 0.27126 | 0.389 |
|  |  | **Aquatic predator surface** | 0.00027 | -0.74525 | 0.758 | 0.00339 | 0.60835 | 0.279 |
|  |  | **Aquatic predator dive** | 0.00015 | -0.94369 | 0.813 | 0.002 | 0.21877 | 0.413 |
|  |  | **Frugivore aerial** | 0.00008 | -1.1111 | 0.858 | 0.00065 | -0.30505 | 0.631 |
|  |  | **Frugivore glean** | 0.0083 | 1.1713 | 0.127 | 0.01503 | 1.80444 | 0.037 * |
|  |  | **Frugivore ground** | 0.00056 | -0.44219 | 0.65 | 0.00062 | -0.40172 | 0.654 |
|  |  | **Nectar aerial** | 0.00339 | 0.6246 | 0.286 | 0.00073 | -0.20786 | 0.593 |
|  |  | **Nectar glean** | 0.00182 | 0.22554 | 0.435 | 0.00359 | 0.65308 | 0.266 |
|  |  | **Granivore above.ground** | 0.00502 | 0.83763 | 0.205 | 0.00057 | -0.42162 | 0.664 |
|  |  | **Granivore ground** | 0.00078 | -0.26161 | 0.613 | 0.00206 | 0.25797 | 0.415 |
|  |  | **Vegetation above.ground** | 0.00058 | -0.40334 | 0.652 | 0.00017 | -0.93151 | 0.814 |
|  |  | **Vegetation ground** | 0.01604 | 1.7118 | 0.042 * | 0.01588 | 1.73519 | 0.04 * |
|  |  | **Vegetation aquatic** | 0.01376 | 1.5815 | 0.046 * | 0.01778 | 1.92995 | 0.025 * |
|  |  | **Vertebrate aerial screening** | 0.00631 | 1.1008 | 0.132 | 0.01563 | 1.68411 | 0.034 * |
|  |  | **Vertebrate aerial to substrate** | 0.00181 | 0.24835 | 0.409 | 0.00001 | -1.68629 | 0.944 |
|  |  | **Vertebrate sally to substrate** | 0.00128 | -0.063041 | 0.533 | 0.00038 | -0.64576 | 0.738 |
|  |  | **Vertebrate glean arboreal** | 0.01313 | 1.5501 | 0.052 . | 0.01266 | 1.57068 | 0.058 . |
|  |  | **Vertebrate glean ground** | 0.00204 | 0.22046 | 0.44 | 0.00442 | 0.70847 | 0.268 |
|  |  | **Scavenger aquatic** | 0.00247 | 0.42854 | 0.344 | 0.00003 | -1.51779 | 0.92 |
|  |  | **Scavenger ground** | 0.00577 | 0.91236 | 0.181 | 0.00633 | 0.98033 | 0.174 |
| **Optic lobe**  K_mult_ = 0.53 | **Shape** | **Invertebrate aerial screen** | 0.01142 | 2.2610 | 0.019 * | 0.00381 | 0.8451 | 0.194 |
|  |  | **Invertebrate aerial sally** | 0.00618 | 1.6233 | 0.060 . | 0.00532 | 1.5745 | 0.058 . |
|  |  | **Invertebrate sally to substrate** | 0.00584 | 1.4538 | 0.073 . | 0.0036 | 0.7324 | 0.238 |
|  |  | **Invertebrate sally to ground** | 0.00190 | -0.5880 | 0.725 | 0.00174 | -0.7801 | 0.79 |
|  |  | **Invertebrate glean bark.rock** | 0.00305 | 0.2777 | 0.385 | 0.0021 | -0.3756 | 0.644 |
|  |  | **Invertebrate glean arboreal** | 0.00449 | 1.0246 | 0.147 | 0.00392 | 0.9254 | 0.18 |
|  |  | **Invertebrate glean ground** | 0.00505 | 1.2321 | 0.104 | 0.00207 | -0.5151 | 0.717 |
|  |  | **Invertebrate aquatic ground** | 0.00954 | 2.0865 | 0.022 * | 0.00599 | 1.6882 | 0.046 * |
|  |  | **Aquatic predator ground** | 0.01794 | 2.5383 | 0.010 ** | 0.0079 | 2.2135 | 0.013 * |
|  |  | **Aquatic predator air** | 0.00935 | 2.1593 | 0.015 * | 0.00129 | -1.3851 | 0.917 |
|  |  | **Aquatic predator plunge** | 0.00488 | 1.0508 | 0.144 | 0.00232 | -0.1961 | 0.57 |
|  |  | **Aquatic predator surface** | 0.00428 | 0.8896 | 0.191 | 0.00257 | 0.1205 | 0.447 |
|  |  | **Aquatic predator dive** | 0.01307 | 2.8235 | 0.003 ** | 0.00893 | 2.3608 | 0.008 ** |
|  |  | **Frugivore aerial** | 0.00208 | -0.3426 | 0.642 | 0.00221 | -0.1063 | 0.546 |
|  |  | **Frugivore glean** | 0.00359 | 0.4933 | 0.305 | 0.00189 | -0.7042 | 0.758 |
|  |  | **Frugivore ground** | 0.00598 | 1.5823 | 0.057 . | 0.00394 | 0.9191 | 0.178 |
|  |  | **Nectar aerial** | 0.00262 | 0.1822 | 0.445 | 0.00279 | 0.3045 | 0.383 |
|  |  | **Nectar glean** | 0.00701 | 1.7197 | 0.045 * | 0.00503 | 1.3595 | 0.094 . |
|  |  | **Granivore above.ground** | 0.00412 | 0.7707 | 0.228 | 0.00283 | 0.1916 | 0.419 |
|  |  | **Granivore ground** | 0.00639 | 1.7871 | 0.037 * | 0.0035 | 0.7193 | 0.24 |
|  |  | **Vegetation above.ground** | 0.00319 | 0.3081 | 0.379 | 0.00283 | 0.2321 | 0.387 |
|  |  | **Vegetation ground** | 0.00727 | 1.9541 | 0.023 * | 0.00753 | 2.1724 | 0.02 * |
|  |  | **Vegetation aquatic** | 0.00237 | -0.1742 | 0.565 | 0.00453 | 1.2488 | 0.102 |
|  |  | **Vertebrate aerial screening** | 0.04597 | 3.1286 | 0.001 *** | 0.00594 | 1.5354 | 0.071 . |
|  |  | **Vertebrate aerial to substrate** | 0.01885 | 2.7775 | 0.002 ** | 0.00538 | 1.472 | 0.063 . |
|  |  | **Vertebrate sally to substrate** | 0.01354 | 2.6219 | 0.006 ** | 0.00393 | 0.9086 | 0.175 |
|  |  | **Vertebrate glean arboreal** | 0.00302 | 0.2271 | 0.402 | 0.0027 | 0.1739 | 0.431 |
|  |  | **Vertebrate glean ground** | 0.00383 | 0.6790 | 0.259 | 0.00386 | 0.8921 | 0.18 |
|  |  | **Scavenger aquatic** | 0.00604 | 1.3453 | 0.098 . | 0.0071 | 1.8798 | 0.032 * |
|  |  | **Scavenger ground** | 0.00933 | 2.2218 | 0.011 * | 0.00475 | 1.2568 | 0.101 |
|  | **Size** | **Invertebrate aerial screen** | 0.00517 | 0.81013 | 0.214 | 0.00203 | 0.29133 | 0.392 |
|  |  | **Invertebrate aerial sally** | 0.00095 | -0.25301 | 0.603 | 0.00389 | 0.64576 | 0.269 |
|  |  | **Invertebrate sally to substrate** | 0.01802 | 1.8242 | 0.029 * | 0.00176 | 0.15459 | 0.467 |
|  |  | **Invertebrate sally to ground** | 0.00213 | 0.22096 | 0.427 | 0.0004 | -0.57687 | 0.694 |
|  |  | **Invertebrate glean bark.rock** | 0.01297 | 1.6688 | 0.043 * | 0.00902 | 1.42273 | 0.075 . |
|  |  | **Invertebrate glean arboreal** | 0.00015 | -1.1122 | 0.846 | 0.00118 | -0.11276 | 0.562 |
|  |  | **Invertebrate glean ground** | 0.03509 | 2.5644 | 0.001 *** | 0.02742 | 2.41654 | 0.005 ** |
|  |  | **Invertebrate aquatic ground** | 0.00423 | 0.7084 | 0.25 | 0.00003 | -1.48416 | 0.918 |
|  |  | **Aquatic predator ground** | 0.00023 | -0.81805 | 0.781 | 0.00014 | -0.9843 | 0.818 |
|  |  | **Aquatic predator air** | 0.01718 | 1.8643 | 0.028 * | 0.0064 | 1.12271 | 0.137 |
|  |  | **Aquatic predator plunge** | 0.00302 | 0.42908 | 0.345 | 0.00185 | 0.2294 | 0.418 |
|  |  | **Aquatic predator surface** | 0.0089 | 1.2866 | 0.104 | 0.00126 | -0.02176 | 0.523 |
|  |  | **Aquatic predator dive** | 0.00769 | 1.194 | 0.118 | 0.00812 | 1.33266 | 0.086 . |
|  |  | **Frugivore aerial** | 0.00058 | -0.41381 | 0.669 | 0.00034 | -0.53728 | 0.706 |
|  |  | **Frugivore glean** | 0.00453 | 0.67955 | 0.265 | 0.00783 | 1.20509 | 0.121 |
|  |  | **Frugivore ground** | 0.01746 | 1.8862 | 0.022 * | 0.01855 | 2.0683 | 0.011 * |
|  |  | **Nectar aerial** | 0.00097 | -0.12505 | 0.555 | 0.00363 | 0.70156 | 0.254 |
|  |  | **Nectar glean** | 0.00044 | -0.59749 | 0.714 | 0.00002 | -1.6587 | 0.944 |
|  |  | **Granivore above.ground** | 0.00081 | -0.28729 | 0.602 | 0.00417 | 0.81449 | 0.219 |
|  |  | **Granivore ground** | 0.00261 | 0.29034 | 0.401 | 0 | -2.24495 | 0.995 |
|  |  | **Vegetation above.ground** | 0.00145 | 0.042895 | 0.485 | 0.00042 | -0.50707 | 0.689 |
|  |  | **Vegetation ground** | 0.00684 | 1.0412 | 0.155 | 0 | -2.10456 | 0.984 |
|  |  | **Vegetation aquatic** | 0.03904 | 2.6379 | 0.004 ** | 0.0413 | 2.81012 | 0.004 ** |
|  |  | **Vertebrate aerial screening** | 0.00322 | 0.59547 | 0.278 | 0.00255 | 0.48289 | 0.322 |
|  |  | **Vertebrate aerial to substrate** | 0.0073 | 1.1014 | 0.137 | 0.00185 | 0.21619 | 0.436 |
|  |  | **Vertebrate sally to substrate** | 0.00956 | 1.3227 | 0.097 . | 0.00372 | 0.75617 | 0.256 |
|  |  | **Vertebrate glean arboreal** | 0.01527 | 1.7573 | 0.035 * | 0.00802 | 1.28928 | 0.104 |
|  |  | **Vertebrate glean ground** | 0.0133 | 1.653 | 0.041 * | 0.00283 | 0.46618 | 0.343 |
|  |  | **Scavenger aquatic** | 0.00432 | 0.76465 | 0.225 | 0.01254 | 1.6758 | 0.04 * |
|  |  | **Scavenger ground** | 0.04054 | 2.7635 | 0.002 ** | 0.01775 | 1.99192 | 0.022 * |
|  | **Relative size** | **Invertebrate aerial screen** | 0.00744 | 1.1363 | 0.144 | 0.0006 | -0.2619 | 0.609 |
|  |  | **Invertebrate aerial sally** | 0.00192 | 0.14067 | 0.46 | 0.02854 | 2.5103 | 0.005 ** |
|  |  | **Invertebrate sally to substrate** | 0.01893 | 1.9562 | 0.019 * | 0.00012 | -1.006 | 0.832 |
|  |  | **Invertebrate sally to ground** | 0.00261 | 0.38927 | 0.365 | 0.00241 | 0.4862 | 0.319 |
|  |  | **Invertebrate glean bark.rock** | 0.00945 | 1.3418 | 0.092 . | 0.00963 | 1.5466 | 0.057 . |
|  |  | **Invertebrate glean arboreal** | 0.00123 | -0.10461 | 0.559 | 0.0003 | -0.6798 | 0.736 |
|  |  | **Invertebrate glean ground** | 0.05072 | 3.1345 | 0.001 *** | 0.05782 | 3.3497 | 0.001 *** |
|  |  | **Invertebrate aquatic ground** | 0.00841 | 1.2178 | 0.111 | 0.00578 | 1.1403 | 0.135 |
|  |  | **Aquatic predator ground** | 0.0038 | 0.6427 | 0.278 | 0.00036 | -0.5138 | 0.686 |
|  |  | **Aquatic predator air** | 0.05195 | 2.9529 | 0.001 *** | 0.00179 | 0.3038 | 0.399 |
|  |  | **Aquatic predator plunge** | 0.00073 | -0.34026 | 0.638 | 0.00263 | 0.5061 | 0.333 |
|  |  | **Aquatic predator surface** | 0.00831 | 1.2174 | 0.108 | 0.00012 | -1 | 0.816 |
|  |  | **Aquatic predator dive** | 0.00851 | 1.2844 | 0.094 . | 0.00131 | 0.1292 | 0.455 |
|  |  | **Frugivore aerial** | 0.0002 | -0.85261 | 0.793 | 0.00009 | -1.0988 | 0.851 |
|  |  | **Frugivore glean** | 0.00293 | 0.38481 | 0.374 | 0.00083 | -0.1992 | 0.592 |
|  |  | **Frugivore ground** | 0.00066 | -0.39879 | 0.656 | 0.00535 | 1.065 | 0.149 |
|  |  | **Nectar aerial** | 0.0001 | -1.0632 | 0.845 | 0.00019 | -0.7855 | 0.773 |
|  |  | **Nectar glean** | 0.00174 | 0.065313 | 0.494 | 0 | -2.4747 | 0.997 |
|  |  | **Granivore above.ground** | 0.00505 | 0.76592 | 0.247 | 0.00729 | 1.3296 | 0.082 . |
|  |  | **Granivore ground** | 0.00945 | 1.2525 | 0.111 | 0.00025 | -0.7835 | 0.768 |
|  |  | **Vegetation above.ground** | 0.00318 | 0.5323 | 0.313 | 0.0015 | 0.218 | 0.432 |
|  |  | **Vegetation ground** | 0.0039 | 0.61469 | 0.283 | 0.00013 | -0.9786 | 0.808 |
|  |  | **Vegetation aquatic** | 0.04043 | 2.7028 | 0.002 ** | 0.03372 | 2.7504 | 0.003 ** |
|  |  | **Vertebrate aerial screening** | 0.05114 | 2.7405 | 0.004 ** | 0.0001 | -1.0508 | 0.841 |
|  |  | **Vertebrate aerial to substrate** | 0.00013 | -1.0135 | 0.842 | 0.00006 | -1.2814 | 0.892 |
|  |  | **Vertebrate sally to substrate** | 0.05404 | 2.9495 | 0.001 *** | 0.03148 | 2.8615 | 0.002 ** |
|  |  | **Vertebrate glean arboreal** | 0.00588 | 0.92072 | 0.185 | 0.00145 | 0.1353 | 0.459 |
|  |  | **Vertebrate glean ground** | 0.00593 | 0.96204 | 0.174 | 0.00752 | 1.2963 | 0.102 |
|  |  | **Scavenger aquatic** | 0.00969 | 1.3713 | 0.088 . | 0.01323 | 1.9246 | 0.022 * |
|  |  | **Scavenger ground** | 0.02664 | 2.3185 | 0.007 ** | 0.01033 | 1.6228 | 0.048 * |
| **Cerebellum**  K_mult_ = 0.14 | **Shape** | **Invertebrate aerial screen** | 0.00369 | 0.62627 | 0.271 | 0.00368 | 0.6491 | 0.267 |
|  |  | **Invertebrate aerial sally** | 0.00532 | 1.1229 | 0.136 | 0.01423 | 2.9558 | 0.003 ** |
|  |  | **Invertebrate sally to substrate** | 0.00223 | -0.15705 | 0.542 | 0.00404 | 0.7593 | 0.232 |
|  |  | **Invertebrate sally to ground** | 0.00106 | -1.4191 | 0.928 | 0.00125 | -1.3305 | 0.902 |
|  |  | **Invertebrate glean bark.rock** | 0.00237 | -0.09524 | 0.534 | 0.00175 | -0.7125 | 0.769 |
|  |  | **Invertebrate glean arboreal** | 0.00117 | -1.6202 | 0.955 | 0.00339 | 0.5035 | 0.315 |
|  |  | **Invertebrate glean ground** | 0.00561 | 1.2107 | 0.120 | 0.00905 | 2.3325 | 0.012 * |
|  |  | **Invertebrate aquatic ground** | 0.04942 | 3.3891 | 0.001 *** | 0.01528 | 3.1809 | 0.001 *** |
|  |  | **Aquatic predator ground** | 0.00279 | 0.38207 | 0.343 | 0.00214 | -0.257 | 0.59 |
|  |  | **Aquatic predator air** | 0.00401 | 0.68918 | 0.233 | 0.00159 | -0.758 | 0.784 |
|  |  | **Aquatic predator plunge** | 0.00258 | 0.15233 | 0.449 | 0.00156 | -0.7979 | 0.77 |
|  |  | **Aquatic predator surface** | 0.00852 | 1.6906 | 0.052 . | 0.00299 | 0.3697 | 0.346 |
|  |  | **Aquatic predator dive** | 0.00473 | 0.8660 | 0.197 | 0.01604 | 2.7145 | 0.001 *** |
|  |  | **Frugivore aerial** | 0.00167 | -0.65324 | 0.752 | 0.00258 | 0.1384 | 0.447 |
|  |  | **Frugivore glean** | 0.00671 | 1.5322 | 0.071 . | 0.0072 | 1.8277 | 0.038 * |
|  |  | **Frugivore ground** | 0.02676 | 3.2156 | 0.001 *** | 0.01498 | 2.9156 | 0.002 ** |
|  |  | **Nectar aerial** | 0.00154 | -0.50855 | 0.694 | 0.00128 | -0.922 | 0.827 |
|  |  | **Nectar glean** | 0.00466 | 0.90486 | 0.194 | 0.00474 | 1.081 | 0.148 |
|  |  | **Granivore above.ground** | 0.00285 | 0.09144 | 0.469 | 0.00651 | 1.6504 | 0.053 . |
|  |  | **Granivore ground** | 0.00231 | -0.31546 | 0.614 | 0.00402 | 0.8384 | 0.2 |
|  |  | **Vegetation above.ground** | 0.00166 | -0.83827 | 0.796 | 0.00281 | 0.1354 | 0.452 |
|  |  | **Vegetation ground** | 0.00175 | -0.68179 | 0.762 | 0.00383 | 0.7583 | 0.232 |
|  |  | **Vegetation aquatic** | 0.00263 | -0.02446 | 0.501 | 0.00535 | 1.2683 | 0.11 |
|  |  | **Vertebrate aerial screening** | 0.01132 | 1.8651 | 0.027 * | 0.00312 | 0.4264 | 0.336 |
|  |  | **Vertebrate aerial to substrate** | 0.00306 | 0.36194 | 0.359 | 0.00157 | -0.8188 | 0.801 |
|  |  | **Vertebrate sally to substrate** | 0.00249 | 0.00064 | 0.493 | 0.00306 | 0.2709 | 0.398 |
|  |  | **Vertebrate glean arboreal** | 0.00215 | -0.25427 | 0.588 | 0.00261 | 0.015 | 0.493 |
|  |  | **Vertebrate glean ground** | 0.00157 | -1.0704 | 0.859 | 0.00279 | 0.1337 | 0.443 |
|  |  | **Scavenger aquatic** | 0.00281 | 0.24277 | 0.394 | 0.00204 | -0.3507 | 0.644 |
|  |  | **Scavenger ground** | 0.00282 | 0.16645 | 0.434 | 0.00271 | 0.1215 | 0.459 |
|  | **Size** | **Invertebrate aerial screen** | 0.00152 | 0.071213 | 0.487 | 0.00224 | 0.33409 | 0.382 |
|  |  | **Invertebrate aerial sally** | 0.00209 | 0.21963 | 0.427 | 0.01175 | 1.62357 | 0.053 . |
|  |  | **Invertebrate sally to substrate** | 0.00064 | -0.41236 | 0.673 | 0.00422 | 0.78442 | 0.234 |
|  |  | **Invertebrate sally to ground** | 0.00026 | -0.7273 | 0.743 | 0.00248 | 0.35429 | 0.374 |
|  |  | **Invertebrate glean bark.rock** | 0.0015 | 0.011682 | 0.508 | 0.00184 | 0.22755 | 0.431 |
|  |  | **Invertebrate glean arboreal** | 0.00014 | -1.027 | 0.83 | 0.00028 | -0.79857 | 0.781 |
|  |  | **Invertebrate glean ground** | 0.02343 | 2.1424 | 0.012 * | 0.02405 | 2.28601 | 0.005 ** |
|  |  | **Invertebrate aquatic ground** | 0.06172 | 3.0857 | 0.002 ** | 0.02248 | 2.15898 | 0.012 * |
|  |  | **Aquatic predator ground** | 0.00114 | -0.021501 | 0.511 | 0.00024 | -0.76556 | 0.772 |
|  |  | **Aquatic predator air** | 0.02201 | 2.0476 | 0.014 * | 0.00003 | -1.43149 | 0.909 |
|  |  | **Aquatic predator plunge** | 0.00275 | 0.46848 | 0.333 | 0.00007 | -1.17954 | 0.87 |
|  |  | **Aquatic predator surface** | 0.00016 | -0.93913 | 0.809 | 0.00172 | 0.17644 | 0.437 |
|  |  | **Aquatic predator dive** | 0.00067 | -0.41242 | 0.667 | 0.00156 | 0.11068 | 0.483 |
|  |  | **Frugivore aerial** | 0.00024 | -0.81802 | 0.786 | 0.00257 | 0.41822 | 0.349 |
|  |  | **Frugivore glean** | 0.00013 | -1.1513 | 0.862 | 0.00128 | -0.02225 | 0.513 |
|  |  | **Frugivore ground** | 0.01684 | 1.8387 | 0.027 * | 0.00501 | 0.90961 | 0.19 |
|  |  | **Nectar aerial** | 0.0009 | -0.19253 | 0.583 | 0.00042 | -0.50988 | 0.695 |
|  |  | **Nectar glean** | 0.00081 | -0.26493 | 0.607 | 0.00006 | -1.23882 | 0.884 |
|  |  | **Granivore above.ground** | 0.00687 | 1.0512 | 0.163 | 0.00818 | 1.35464 | 0.096 |
|  |  | **Granivore ground** | 0.00815 | 1.1986 | 0.12 | 0 | -2.25443 | 0.992 |
|  |  | **Vegetation above.ground** | 0.00063 | -0.312 | 0.636 | 0.00037 | -0.62704 | 0.732 |
|  |  | **Vegetation ground** | 0.00092 | -0.17642 | 0.583 | 0.00121 | -0.0713 | 0.534 |
|  |  | **Vegetation aquatic** | 0.00181 | 0.12893 | 0.472 | 0.00288 | 0.5338 | 0.321 |
|  |  | **Vertebrate aerial screening** | 0.02238 | 2.0987 | 0.015 * | 0.02047 | 2.03127 | 0.012 * |
|  |  | **Vertebrate aerial to substrate** | 0.00004 | -1.4787 | 0.92 | 0.00131 | 0.02103 | 0.493 |
|  |  | **Vertebrate sally to substrate** | 0.0142 | 1.697 | 0.033 * | 0.01785 | 1.94209 | 0.022 * |
|  |  | **Vertebrate glean arboreal** | 0.00138 | -0.050898 | 0.533 | 0.00206 | 0.26029 | 0.416 |
|  |  | **Vertebrate glean ground** | 0.00193 | 0.146 | 0.459 | 0.00362 | 0.6317 | 0.287 |
|  |  | **Scavenger aquatic** | 0.0013 | 0.039382 | 0.491 | 0.00001 | -1.71641 | 0.955 |
|  |  | **Scavenger ground** | 0.00001 | -1.804 | 0.953 | 0.00001 | -1.90421 | 0.968 |
|  | **Relative size** | **Invertebrate aerial screen** | 0.00201 | 0.2434 | 0.421 | 0.0036 | 0.6582 | 0.265 |
|  |  | **Invertebrate aerial sally** | 0.00406 | 0.6619 | 0.262 | 0.00593 | 1.02593 | 0.152 |
|  |  | **Invertebrate sally to substrate** | 0.00059 | -0.43808 | 0.682 | 0.00254 | 0.39952 | 0.373 |
|  |  | **Invertebrate sally to ground** | 0.00041 | -0.52447 | 0.685 | 0.00269 | 0.39177 | 0.363 |
|  |  | **Invertebrate glean bark.rock** | 0.0012 | -0.1057 | 0.556 | 0.00104 | -0.1031 | 0.544 |
|  |  | **Invertebrate glean arboreal** | 0.00003 | -1.5208 | 0.926 | 0.00026 | -0.83189 | 0.795 |
|  |  | **Invertebrate glean ground** | 0.02372 | 2.1451 | 0.011 * | 0.02205 | 2.12027 | 0.01 ** |
|  |  | **Invertebrate aquatic ground** | 0.05194 | 2.8986 | 0.002 ** | 0.02222 | 2.11238 | 0.015 * |
|  |  | **Aquatic predator ground** | 0.0001 | -1.0845 | 0.85 | 0.00021 | -0.8241 | 0.793 |
|  |  | **Aquatic predator air** | 0.01512 | 1.7081 | 0.043 * | 0.0003 | -0.69642 | 0.746 |
|  |  | **Aquatic predator plunge** | 0.00316 | 0.55636 | 0.307 | 0 | -2.43657 | 0.996 |
|  |  | **Aquatic predator surface** | 0.00051 | -0.44007 | 0.656 | 0.00117 | -0.06249 | 0.532 |
|  |  | **Aquatic predator dive** | 0.00074 | -0.37403 | 0.646 | 0.00076 | -0.27197 | 0.616 |
|  |  | **Frugivore aerial** | 0.00058 | -0.44528 | 0.667 | 0.0024 | 0.34698 | 0.382 |
|  |  | **Frugivore glean** | 0.00032 | -0.83442 | 0.775 | 0.00079 | -0.2999 | 0.62 |
|  |  | **Frugivore ground** | 0.01144 | 1.4668 | 0.07 . | 0.00375 | 0.6666 | 0.271 |
|  |  | **Nectar aerial** | 0 | -2.3003 | 0.992 | 0.00002 | -1.5598 | 0.93 |
|  |  | **Nectar glean** | 0.00071 | -0.33804 | 0.634 | 0.00022 | -0.83783 | 0.795 |
|  |  | **Granivore above.ground** | 0.00733 | 1.099 | 0.142 | 0.00998 | 1.50862 | 0.067 |
|  |  | **Granivore ground** | 0.00659 | 1.0314 | 0.161 | 0.00004 | -1.44734 | 0.907 |
|  |  | **Vegetation above.ground** | 0.00038 | -0.57507 | 0.707 | 0.00016 | -0.97618 | 0.822 |
|  |  | **Vegetation ground** | 0.00116 | -0.060397 | 0.54 | 0.00204 | 0.23671 | 0.415 |
|  |  | **Vegetation aquatic** | 0.00262 | 0.34723 | 0.375 | 0.00299 | 0.54435 | 0.319 |
|  |  | **Vertebrate aerial screening** | 0.00805 | 1.2529 | 0.097 . | 0.01994 | 1.96005 | 0.015 * |
|  |  | **Vertebrate aerial to substrate** | 0.0017 | 0.11675 | 0.473 | 0.00293 | 0.51557 | 0.325 |
|  |  | **Vertebrate sally to substrate** | 0.00419 | 0.74366 | 0.247 | 0.00987 | 1.41561 | 0.079 . |
|  |  | **Vertebrate glean arboreal** | 0.00102 | -0.21599 | 0.585 | 0.00158 | 0.09234 | 0.484 |
|  |  | **Vertebrate glean ground** | 0.0012 | -0.10266 | 0.557 | 0.00405 | 0.68617 | 0.253 |
|  |  | **Scavenger aquatic** | 0.00173 | 0.20644 | 0.428 | 0.00011 | -1.064 | 0.828 |
|  |  | **Scavenger ground** | 0.0002 | -0.88928 | 0.81 | 0.00001 | -1.96139 | 0.97 |
| **Brainstem**  K_mult_ = 0.27 | **Shape** | **Invertebrate aerial screen** | 0.00687 | 1.5241 | 0.067 . | 0.00875 | 2.3862 | 0.008 ** |
|  |  | **Invertebrate aerial sally** | 0.00602 | 1.3865 | 0.084 . | 0.01015 | 2.5704 | 0.002 ** |
|  |  | **Invertebrate sally to substrate** | 0.00457 | 0.9911 | 0.160 | 0.00612 | 1.714 | 0.047 * |
|  |  | **Invertebrate sally to ground** | 0.00279 | 0.1611 | 0.441 | 0.00493 | 1.2456 | 0.108 |
|  |  | **Invertebrate glean bark.rock** | 0.00448 | 0.9575 | 0.178 | 0.00289 | 0.349 | 0.352 |
|  |  | **Invertebrate glean arboreal** | 0.01091 | 2.5192 | 0.007 ** | 0.00536 | 1.5497 | 0.052 . |
|  |  | **Invertebrate glean ground** | 0.00488 | 1.1043 | 0.144 | 0.00351 | 0.7416 | 0.236 |
|  |  | **Invertebrate aquatic ground** | 0.01273 | 2.4231 | 0.011 * | 0.00751 | 2.0309 | 0.023 * |
|  |  | **Aquatic predator ground** | 0.01598 | 2.2973 | 0.014 * | 0.00396 | 0.8929 | 0.178 |
|  |  | **Aquatic predator air** | 0.02655 | 2.9379 | 0.002 ** | 0.00115 | -1.4095 | 0.926 |
|  |  | **Aquatic predator plunge** | 0.00145 | -0.9451 | 0.830 | 0.00124 | -1.1997 | 0.884 |
|  |  | **Aquatic predator surface** | 0.00221 | -0.2825 | 0.598 | 0.00188 | -0.4318 | 0.665 |
|  |  | **Aquatic predator dive** | 0.00807 | 1.8330 | 0.032 * | 0.01446 | 3.1414 | 0.001 *** |
|  |  | **Frugivore aerial** | 0.00162 | -0.7248 | 0.772 | 0.00253 | 0.1712 | 0.433 |
|  |  | **Frugivore glean** | 0.00382 | 0.6566 | 0.256 | 0.00309 | 0.4625 | 0.304 |
|  |  | **Frugivore ground** | 0.00447 | 0.9126 | 0.176 | 0.00506 | 1.3786 | 0.083 . |
|  |  | **Nectar aerial** | 0.00186 | -0.310 | 0.638 | 0.00215 | -0.0895 | 0.551 |
|  |  | **Nectar glean** | 0.00520 | 1.1013 | 0.146 | 0.00332 | 0.5638 | 0.294 |
|  |  | **Granivore above.ground** | 0.00314 | 0.3073 | 0.381 | 0.00264 | 0.1742 | 0.42 |
|  |  | **Granivore ground** | 0.00624 | 1.4682 | 0.069 . | 0.00528 | 1.426 | 0.08 . |
|  |  | **Vegetation above.ground** | 0.00158 | -0.9814 | 0.840 | 0.00174 | -0.6762 | 0.752 |
|  |  | **Vegetation ground** | 0.00283 | 0.0478 | 0.482 | 0.00422 | 1.0223 | 0.153 |
|  |  | **Vegetation aquatic** | 0.00345 | 0.4965 | 0.321 | 0.00289 | 0.4317 | 0.32 |
|  |  | **Vertebrate aerial screening** | 0.06876 | 3.1558 | 0.001 *** | 0.00513 | 1.2593 | 0.112 |
|  |  | **Vertebrate aerial to substrate** | 0.04434 | 3.4649 | 0.001 *** | 0.0159 | 3.6219 | 0.001 *** |
|  |  | **Vertebrate sally to substrate** | 0.02028 | 2.9985 | 0.004 ** | 0.00977 | 2.5991 | 0.008 ** |
|  |  | **Vertebrate glean arboreal** | 0.00331 | 0.4294 | 0.327 | 0.00148 | -0.9696 | 0.834 |
|  |  | **Vertebrate glean ground** | 0.00266 | -0.0528 | 0.517 | 0.00223 | -0.1654 | 0.558 |
|  |  | **Scavenger aquatic** | 0.00279 | 0.1603 | 0.438 | 0.00378 | 0.7621 | 0.224 |
|  |  | **Scavenger ground** | 0.00633 | 1.3757 | 0.088 . | 0.00408 | 0.9712 | 0.176 |
|  | **Size** | **Invertebrate aerial screen** | 0.01706 | 1.783 | 0.033 * | 0.0041 | 0.7905 | 0.226 |
|  |  | **Invertebrate aerial sally** | 0.00185 | 0.1718 | 0.438 | 0.02769 | 2.55298 | 0.001 *** |
|  |  | **Invertebrate sally to substrate** | 0.00357 | 0.58751 | 0.286 | 0.00037 | -0.71216 | 0.754 |
|  |  | **Invertebrate sally to ground** | 0.00001 | -1.7447 | 0.961 | 0.0028 | 0.60951 | 0.293 |
|  |  | **Invertebrate glean bark.rock** | 0.00458 | 0.73337 | 0.251 | 0.00072 | -0.26599 | 0.593 |
|  |  | **Invertebrate glean arboreal** | 0.00027 | -0.75698 | 0.766 | 0.00022 | -0.80758 | 0.775 |
|  |  | **Invertebrate glean ground** | 0.00029 | -0.83316 | 0.792 | 0.0114 | 1.59637 | 0.053 . |
|  |  | **Invertebrate aquatic ground** | 0.00022 | -0.81198 | 0.779 | 0.00003 | -1.41799 | 0.906 |
|  |  | **Aquatic predator ground** | 0.02014 | 1.943 | 0.025 * | 0.00086 | -0.2109 | 0.586 |
|  |  | **Aquatic predator air** | 0.02789 | 2.3609 | 0.007 ** | 0.00497 | 0.87944 | 0.214 |
|  |  | **Aquatic predator plunge** | 0.00132 | 0.0018971 | 0.526 | 0.00007 | -1.16162 | 0.855 |
|  |  | **Aquatic predator surface** | 0.0002 | -0.87875 | 0.805 | 0.00511 | 0.94346 | 0.18 |
|  |  | **Aquatic predator dive** | 0.00066 | -0.41009 | 0.652 | 0.00091 | -0.15984 | 0.568 |
|  |  | **Frugivore aerial** | 0.0011 | -0.068793 | 0.533 | 0.00006 | -1.24236 | 0.873 |
|  |  | **Frugivore glean** | 0.00132 | -0.097433 | 0.553 | 0.01926 | 2.05972 | 0.013 * |
|  |  | **Frugivore ground** | 0.00386 | 0.62095 | 0.284 | 0.00646 | 1.1321 | 0.135 |
|  |  | **Nectar aerial** | 0.00001 | -1.6581 | 0.942 | 0.00062 | -0.26656 | 0.615 |
|  |  | **Nectar glean** | 0.00551 | 0.872 | 0.212 | 0.00188 | 0.19446 | 0.448 |
|  |  | **Granivore above.ground** | 0.00279 | 0.4298 | 0.348 | 0.00241 | 0.44428 | 0.358 |
|  |  | **Granivore ground** | 0.00224 | 0.26661 | 0.426 | 0.01353 | 1.80102 | 0.028 * |
|  |  | **Vegetation above.ground** | 0.0079 | 1.2167 | 0.107 | 0.00443 | 0.82568 | 0.219 |
|  |  | **Vegetation ground** | 0.00317 | 0.43108 | 0.342 | 0.00854 | 1.30212 | 0.097 . |
|  |  | **Vegetation aquatic** | 0.01861 | 1.9842 | 0.018 * | 0.00625 | 1.16931 | 0.136 |
|  |  | **Vertebrate aerial screening** | 0.11641 | 3.8592 | 0.001 *** | 0.00009 | -1.15727 | 0.853 |
|  |  | **Vertebrate aerial to substrate** | 0.04772 | 2.7451 | 0.003 ** | 0.01077 | 1.5493 | 0.061 . |
|  |  | **Vertebrate sally to substrate** | 0.05504 | 3.2404 | 0.001 *** | 0.02849 | 2.84999 | 0.002 ** |
|  |  | **Vertebrate glean arboreal** | 0.00446 | 0.73791 | 0.238 | 0.00821 | 1.36566 | 0.084 . |
|  |  | **Vertebrate glean ground** | 0.00125 | -0.078635 | 0.535 | 0.00754 | 1.30863 | 0.099 . |
|  |  | **Scavenger aquatic** | 0.00938 | 1.3308 | 0.085 . | 0.0074 | 1.26344 | 0.104 |
|  |  | **Scavenger ground** | 0.00002 | -1.5002 | 0.92 | 0.00123 | 0.00132 | 0.515 |
|  | **Relative size** | **Invertebrate aerial screen** | 0.01736 | 1.7705 | 0.038 * | 0.00249 | 0.38904 | 0.36 |
|  |  | **Invertebrate aerial sally** | 0.00047 | -0.49294 | 0.694 | 0.01076 | 1.5204 | 0.062 . |
|  |  | **Invertebrate sally to substrate** | 0.00343 | 0.54433 | 0.316 | 0.00025 | -0.90952 | 0.81 |
|  |  | **Invertebrate sally to ground** | 0.00002 | -1.6177 | 0.937 | 0.00322 | 0.65298 | 0.284 |
|  |  | **Invertebrate glean bark.rock** | 0.0063 | 0.94798 | 0.184 | 0.00245 | 0.37128 | 0.377 |
|  |  | **Invertebrate glean arboreal** | 0 | -2.1308 | 0.987 | 0.00048 | -0.5309 | 0.68 |
|  |  | **Invertebrate glean ground** | 0.00148 | -0.036035 | 0.536 | 0.00677 | 1.06875 | 0.149 |
|  |  | **Invertebrate aquatic ground** | 0.00452 | 0.76861 | 0.227 | 0.00114 | -0.05745 | 0.534 |
|  |  | **Aquatic predator ground** | 0.00855 | 1.2763 | 0.088 . | 0.00029 | -0.73952 | 0.759 |
|  |  | **Aquatic predator air** | 0.01584 | 1.8043 | 0.031 * | 0.00329 | 0.5035 | 0.337 |
|  |  | **Aquatic predator plunge** | 0.00131 | 0.0083149 | 0.513 | 0.00001 | -1.62167 | 0.936 |
|  |  | **Aquatic predator surface** | 0.00088 | -0.2693 | 0.603 | 0.00326 | 0.57839 | 0.308 |
|  |  | **Aquatic predator dive** | 0.00067 | -0.39158 | 0.64 | 0.00001 | -1.70646 | 0.948 |
|  |  | **Frugivore aerial** | 0.00061 | -0.33626 | 0.633 | 0.0006 | -0.38482 | 0.645 |
|  |  | **Frugivore glean** | 0.00258 | 0.31778 | 0.397 | 0.01808 | 1.90595 | 0.022 * |
|  |  | **Frugivore ground** | 0.01858 | 1.9854 | 0.018 * | 0.01556 | 1.82957 | 0.031 * |
|  |  | **Nectar aerial** | 0.00177 | 0.22602 | 0.425 | 0.00023 | -0.77528 | 0.774 |
|  |  | **Nectar glean** | 0.00512 | 0.78863 | 0.232 | 0.00364 | 0.60997 | 0.283 |
|  |  | **Granivore above.ground** | 0.00186 | 0.16881 | 0.448 | 0.00337 | 0.5809 | 0.301 |
|  |  | **Granivore ground** | 0.00045 | -0.58524 | 0.7 | 0.01343 | 1.77174 | 0.033 * |
|  |  | **Vegetation above.ground** | 0.00686 | 1.0859 | 0.136 | 0.00339 | 0.60101 | 0.3 |
|  |  | **Vegetation ground** | 0.0038 | 0.52393 | 0.315 | 0.01369 | 1.62909 | 0.05 * |
|  |  | **Vegetation aquatic** | 0.0193 | 1.9739 | 0.02 * | 0.00865 | 1.37927 | 0.085 . |
|  |  | **Vertebrate aerial screening** | 0.07353 | 3.4222 | 0.002 ** | 0.00031 | -0.7138 | 0.753 |
|  |  | **Vertebrate aerial to substrate** | 0.03126 | 2.2905 | 0.008 ** | 0.00857 | 1.30486 | 0.088 . |
|  |  | **Vertebrate sally to substrate** | 0.02452 | 2.3232 | 0.007 ** | 0.01179 | 1.72732 | 0.039 * |
|  |  | **Vertebrate glean arboreal** | 0.00319 | 0.50144 | 0.338 | 0.00657 | 1.12239 | 0.133 |
|  |  | **Vertebrate glean ground** | 0.00365 | 0.59273 | 0.304 | 0.00807 | 1.30461 | 0.095 . |
|  |  | **Scavenger aquatic** | 0.01161 | 1.5258 | 0.054 . | 0.01139 | 1.58257 | 0.059 . |
|  |  | **Scavenger ground** | 0.00064 | -0.32184 | 0.637 | 0.00072 | -0.28216 | 0.626 |

**Table S3:** List and descriptions of discrete, curve, and surface (semi)landmarks. ‘Discrete landmarks’ column provides anatomical definitions for each discrete landmarks. ‘’Curves’ column lists the discrete landmarks that serve as endpoints along each curve and the number of equidistant semilandmarks sampled along each curve. ‘Surfaces’ column lists curves that bound the region the number of surface semilandmarks placed in each region.

| **Region** | **Discrete landmarks (LM)** | **Curves** | **Surfaces** |
| --- | --- | --- | --- |
| **Cerebrum** | 1. Anterior tip of the cerebrum at the base of the olfactory tract on dorsal side.  2. Posteromedial point of the right cerebrum on dorsal side.  3. Right, dorsal-most junction point of cerebrum and optic lobe.  4. Ventral-most junction point of cerebrum and optic lobe. | 1. LM 1 🡪 2 (10 pts)  2. LM 2 🡪 3 (10 pts)  3. LM 3 🡪 4 (10 pts)  4. LM 4 🡪 1 (15 pts) | Right cerebral hemisphere, within space enclosed by curves 1 – 4 and LM 1 – 4 (52 pts) |
| **Optic Lobe** | 5. Right junction point of optic lobe, midbrain, and brainstem.  6. Right junction of optic lobe, cerebellum, and brainstem. | 5. LM 4 🡪 5 (7 pts)  6. LM 5 🡪 6 (7 pts)  7. LM 6 🡪 3 (7 pts) | Right optic lobe, within space enclosed by curves 5 – 7 and LM 5 –6 (15 pts) |
| **Cerebellum** | 7. Anterior-most median point of cerebellum on dorsal side.  8. Right anteroventral point of the cerebellum.  9. Right posterolateral point of the cerebellum.  10. Posterior-most median point of the cerebellum on dorsal side. | 8. LM 7 🡪 10 (10 pts)  9. LM 10 🡪 9 (5 pts)  10. LM 9 🡪 8 (10 pts)  11. LM 8 🡪7 (10 pts) | Right side of cerebellum, within space enclosed by curves 8 – 11 and LM 7 – 10 (18 pts) |
| **Brainstem** | 11. Anterior-most median point adjacent to midbrain on ventral side.  12. Right posterolateral points of brainstem on ventral surface.  13. Posterior-most median point of brainstem on ventral surface. | 12. LM 11 🡪 13 (10 pts)  13. LM 13 🡪 12 (7 pts)  14. LM 12 🡪 6 (7 pts)  15. LM 5 🡪 11 (5 pts) | Right side of brainstem, within space enclosed by curves 12 – 15 and LM 11– 13 (13 pts) |

**Table S4:** Definitions and Quantifications of Ecomorphological Traits. Traits are organized by their respective ecological axis (Locomotion or Diet) to illustrate the structural differences between binary character coding and continuous proportion indices used as independent predictors in our PGLS regressions. Traits are compiled and adapted from ^4,5^.

| **Ecological Axis** | **Variable Name** | **Data Type** | **Definition** |
| --- | --- | --- | --- |
| **Locomotion** | **Aquatic** | Binary (0/1) | Specialized morphology or behavior for locomotion on or under water. |
|  | **Terrestrial** | Binary (0/1) | Ground-based cursorial locomotion (walking, running, hopping) as a primary foraging or escape mechanism. |
|  | **Insessorial** | Binary (0/1) | Anatomical or behavioral specialization for perching in trees and navigating arboreal vegetation and branches. |
|  | **Soaring** | Binary (0/1) | Morphological adaptations (e.g., high aspect-ratio wings) and behaviors for passive, low-energy thermal (using thermal updrafts) or dynamic (using wind gradients) soaring flight. |
|  | **Volant** | Binary (0/1) | Capacity for sustained, powered aerodynamic flight. A score of 0 denotes complete flightlessness. |
| **Diet** | **Frugivore** | Continuous (0–100%) | Estimated percentage of diet consisting of fleshy fruits |
|  | **Nectarivore** | Continuous (0–100%) | Estimated percentage of diet consisting of floral nectar |
|  | **Granivore** | Continuous (0–100%) | Estimated percentage of diet consisting of seeds and grains |
|  | **Herbivore** | Continuous (0–100%) | Estimated percentage of diet consisting of vegetative plant tissues |
|  | **Invertivore** | Continuous (0–100%) | Estimated percentage of diet consisting of terrestrial or aerial invertebrates (e.g., insects, spiders, worms) |
|  | **Vertivore** | Continuous (0–100%) | Estimated percentage of diet consisting of terrestrial vertebrates (e.g., mammals, birds, reptiles) |
|  | **Aquatic Predator** | Continuous (0–100%) | Estimated percentage of diet consisting of organisms captured in marine or freshwater ecosystems (e.g., fish, aquatic macroinvertebrates) |
|  | **Scavenger** | Continuous (0–100%) | Estimated percentage of diet obtained via the consumption of carrion and animal carcasses. |
|  | **Omnivore** | Discrete category only | The proportional calculation of a multi-resource diet where no single food category constitutes ≥60% of total volumetric intake |
